# Methylation in NDUFA13 gene promoter disrupts communication between collaborative transcription factors – potential mechanism for onset of breast cancer

**DOI:** 10.1101/2022.06.01.494372

**Authors:** Hörberg Johanna, Hallbäck Björn, Moreau Kevin, Anna Reymer

## Abstract

Selective DNA binding by transcription factors (TFs) is crucial for the correct regulation of DNA transcription. In healthy cells, promoters of active genes are hypomethylated. A single CpG methylation within a TF response element may change the binding preferences of the protein thus causing the dysregulation of transcription programs. Here we investigate a molecular mechanism driving the downregulation of NDUFA13 gene, due to hypermethylation, which is associated with multiple cancers. Using bioinformatic analyses of breast cancer cell line MCF7, we identify a hypermethylated region containing the binding sites of two TFs dimers, CEBPB and E2F1-DP1, located 130 b.p. from the gene transcription start site. All-atom extended MD simulations of wild-type and methylated DNA alone and in complex with either one or both TFs dimers provide mechanistic insights into the cooperative asymmetric binding order of the two dimers; the CEBPB binding should occur first to facilitate the E2F1-DP1-DNA association. The CpG methylation within the E2F1-DP1 response element and the linker decreases the cooperativity effects and renders the E2F1-DP1 binding site less recognizable by the TF dimer. Taken together, the identified CpG methylation site may contribute to the downregulation of NDUFA13 gene and has a potential as a biomarker for breast cancer.

## Introduction

DNA methylation is a common epigenetic modification, important for the regulation of neural development, neurogenesis, synaptic plasticity, brain function, immune response, etc (1–3). The genome-wide DNA methylation pattern, established at an early embryonic stage, is maintained through every cell division cycle, though changes can occur as a result of environmental factors or aging (4–6) and may lead to onset of diseases, including cancers (7–10). Changes in DNA methylation pattern can lead to impaired transcription of certain guard genes, and depending on the location whether in gene exons, introns, enhancers or promoters, impact transcription differently (11). The molecular mechanisms for the impact of abnormal DNA methylation on transcriptional regulation vary. For example, increased methylation of CpG steps is associated with heterochromatin (12, 13), which makes the genome more compact and inaccessible for the transcription machinery. Alternatively, the hypermethylation in non-coding genome may impair the binding of regulatory proteins, e.g. transcription factors, or create binding sites for others (14, 15).

DNA methylation is commonly found within a CpG dinucleotide, when a methyl group is added to a C5 atom of two cytosine bases on the opposite DNA strands. There are also instances of a single cytosine methylation, irrespective of the flanking nucleotide. The addition of a methyl group occurs from the major groove and does not affect the Watson-Crick base pairing, instead it modifies the conformational dynamics of DNA, increasing the overall bending and torsional rigidity (16–18). Depending on the local sequence context, the local effects of a CpG methylation site on DNA structure can be significant (19). The local structural changes include increased roll and decreased twist angles (17), and narrowing of the minor groove (20). These structural changes affect DNA-protein interactions; however, the magnitude of the effect depends on the recognition motif and 3D fold of the proteins DNA binding domains. Furthermore, the addition of a methyl group creates additional surfaces for hydrophobic interactions, which can enhance or reduce the protein-DNA binging affinities, though the effect varies depending on where in a response element the modification occurs (21). Thus, understanding the changes cytosine methylation brings into solo and cooperative transcription factors DNA recognition and, by extension, transcriptional regulatory processes is of great importance for the development of early cancer diagnostics, where particular DNA methylation sites can serve as biomarkers (22).

Here we investigate a molecular mechanism driving the downregulation of NDUFA13 gene, due to hypermethylation, which is linked to the onset of multiple cancers (11, 23, 24). We employ bioinformatics analyses of breast cancer cell line MCF7 to derive our model system, based on the promoter of NDUFA13 gene. We identify a hypermethylated region within the promoter of NDUFA13 gene, 130 b.p. from the transcription start site (TSS), which contains the binding sites of two transcription factor (TF) proteins, a homodimer CEBPB and a heterodimer E2F1-DP1. We investigate with all-atom microsecond-long molecular dynamics (MD) simulations the cooperative DNA recognition by the two TF dimers and how the process is affected by the abnormal methylation. The DNA region has six methylation marks, four single methylated cytosines within the binding site of E2F1-DP1 and in the linker, and one double methylation in the flanks of the CEBPB response element. We observe that the methylation reduces the asymmetric DNA-mediated allosteric communication from the CEBPB to E2F1-DP1 binding sites, important for the tighter association of E2F1-DP1 on DNA. Furthermore, the methylation changes the physical properties of the E2F1-DP1-DNA binding site, which results in the loss of specificity in the protein-DNA contacts but has no effect on the CEBPB-DNA binding. Taken together, the identified hypermethylation marks will impede the formation of the CEBPB-E2F1-DP1-DNA enhanceosome complex, and given the close location to the TSS, may significantly contribute to the downregulation of NDUFA13 gene.

## Methods

### Bioinformatic analyses

The studied model system was derived from bioinformatic analyses. We employ a strategy of selecting human promoter regions where abnormal cytosine methylation is present along with transcription factors (TFs) binding sites in the vicinity of transcription starting sites (TSS). Firstly, we filter out regions that possess multiple overlapping ChIP-seq signals from the ENCODE collection for breast cancer cell line MCF7 (25), retaining overlapping ChIP-seq signals within promoter regions (setting the window to 200 b.p. to TSS). Secondly, to further narrow the selection, the methylation pattern in promoters, obtained though Reduced Bisulfite Seq, is analyzed with UCSC Genome Browser with setting ENCODE track to “MCF7 cells results only” (26). After the filtering, the NDUFA13 (GRIM19) promoter (23, 24) is selected. Lastly, we use the TFregulomeR tool (27) to identify TF response elements corresponding to the ChIP-seq signals near the methylated CpG sites.

### Simulated systems

We simulate eight systems, including wild type (WT) and methylated (ME) 49 b.p. long DNA molecules containing two response elements of human TFs dimers: E2F1-DP1 and CEBPB, alone or in complex with either one, or both TFs dimers. The ME-systems DNA contain six methylated cytosines, four single methylation sites within the E2F1-DP1 response element and the linker region, and one double methylation site in the 3’-flanking sequence of the CEBPB response element. See Figure S1 and Figure 1 for WT and ME DNA sequences.

**Figure 1.**
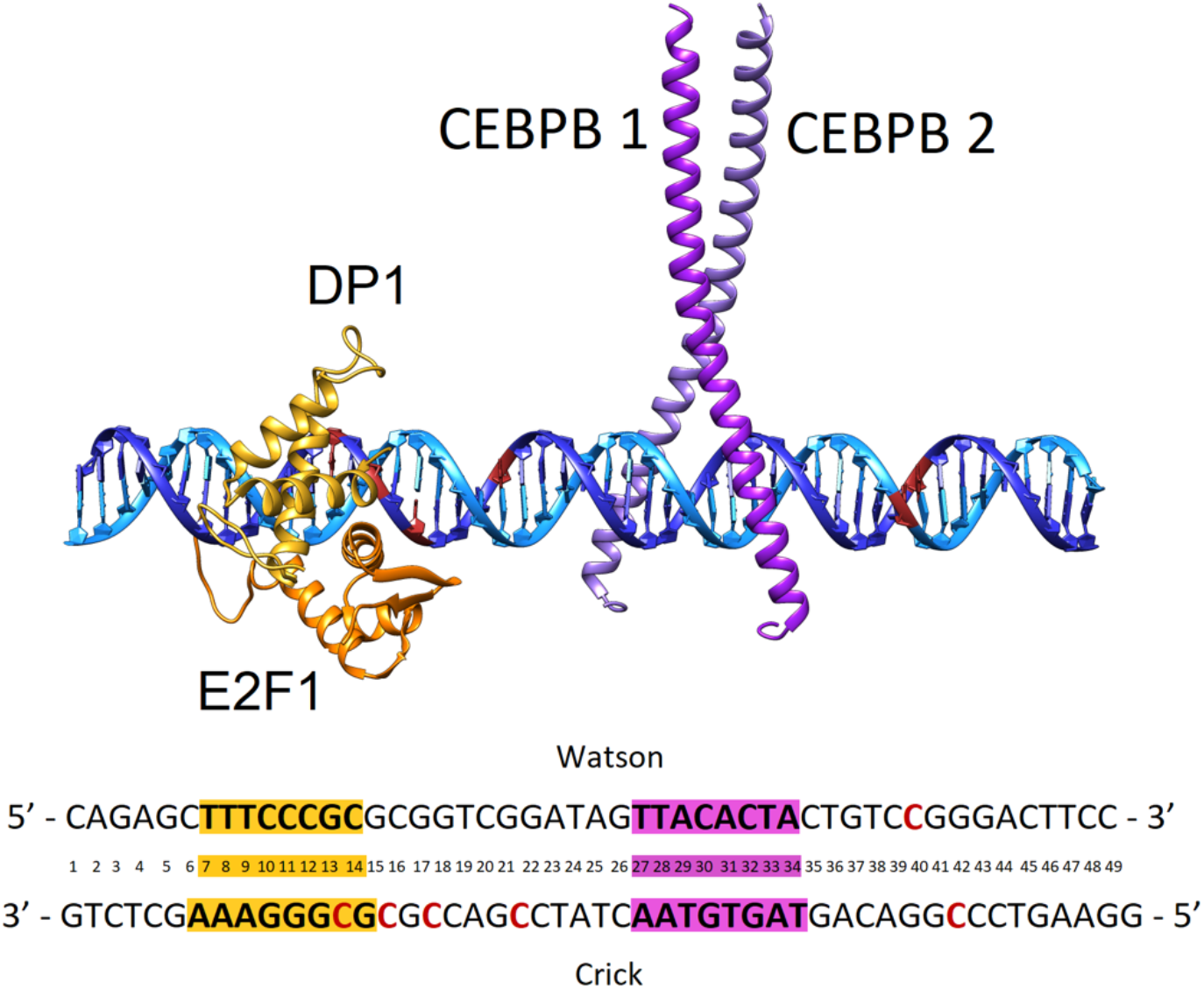
Model system structure of E2F1-DP1-CEBPB-DNA enhanceosome complex. DNA Watson strand (5’->3’ direction) in light blue, DNA Crick strand (3’->5’ direction) in dark blue, with cytosine residues where methylation occurs in red, E2F1 monomer in orange, DP1 monomer in yellow, CEBPB monomer 1 in magenta, and CEBPB monomer 2 in purple. Below is the DNA b.p. numbering used in the model with cytosine methylation sites in red and response elements in yellow and magenta for E2F1-DP1 and CEBPB dimers, respectively.

To design model structures of the systems, we first derive a homology model of the human E2F1-DP1 dimer with YASARA (28, 29) using as the template the crystal structure of the E2F4-DP2-DNA complex (PDB ID: 1CF7 (30)). The sequence identity and sequence similarity (Figure S2A-B) are 63% and 71%, respectively, for the E2F1/E2F4 homologs and 89% and 95%, respectively, for the DP1/DP2 homologs, which justifies the choice of the template and ensures the quality of the model structure. Using HDOCK webserver (31), we proceed with molecular docking of two TFs dimers to B-DNA containing their corresponding response elements and two flanking nucleotides on each side (XXREXX). The structure of the CEBPB dimer is adopted from the crystal structure of the CEBPB-DNA complex (PDB ID: 1GU4). We select the best TF-dimer complex model based on the docking score and similarity to the corresponding crystal structures. Finally, we assemble the corresponding DNA complexes with either one TF dimer, or both in USCF Chimera (32).

### Molecular Dynamics Simulations

All molecular dynamics simulations were performed using the GROMACS MD engine version 2021 (33), using amber14SB (34) and parmbsc1 (35) forcefields for the protein and DNA, respectively, and previously derived parameters for 5-methylcytosine (16, 36). All simulated systems were first neutralised and then solvated with 13Å SPC/E (37) water. Additional K^+^ and Cl^-^ ions were added to reach a physiological concentration of 150mM KCl. Using periodic boundary conditions, the systems were energy minimized with 5000 steps of steepest decent, followed by 500 ps equilibration-runs with weak position restraints on heavy atoms of the solute (1000 kJ/mol) in the NVT and NPT ensembles, adjusting temperature and pressure to 300 K and 1 atm (38, 39). Releasing the restraints, for each of eight systems we carry out 1.1 µs MD simulations at constant pressure and temperature (1 atm and 300 K).

### Trajectory Analyses

For each of the generated MD trajectories the first 100 ns are discarded as equilibration. CPPTRAJ program from AMBERTOOLS 16 software package (40) is used for the analysis of protein-DNA contacts. We analyse both specific (hydrogen bonding and hydrophobic contacts) and nonspecific contacts (formed between either DNA or protein backbones) that are present for longer than 10% of the trajectory, see previous publication for details (41). Subsequently, Curves+, Canal, and Canion programs (42) are used to derive the helical parameters, backbone torsional angles, groove geometry parameters, and ion distributions for each trajectory snapshot extracted every ps. GROMACS energy tool is used to calculate protein-DNA interaction energies, that include short range electrostatic and Lennard-Jones interactions. To separate interaction energies into specific and nonspecific, we calculate interactions for several atom groups that correspond to DNA bases, protein side chains, and molecule backbone. Analysis of free energies is performed using the MMPBSA/MMGBSA plugin in AMBERTOOLS 16. DNA deformation energies were derived with a multivariate Ising model (43), which combines a harmonic deformation approximation model with an Ising model to allow for inclusion of coupling between all conformational substates of DNA. The model has been parameterized for all tetranucleotides. The model utilizes six inter-base pair (shift, slide, rise, twist, tilt, and roll) and the six intra-base pair (shear, stagger, stretch, buckle, propeller twist, and opening) parameters to calculate the deformation energy for a DNA sequence. Long-distance correlation analysis between DNA translational helical and groove parameters was performed in MatLab software, according to the methods of Balaceanu et al. (44).

### Additional Information

MatLab software and Microsoft excel were used for the post-processing and plotting of all data. USCF Chimera (32) was used for creating all molecular graphics.

## Results

Using computational methods, we investigate a molecular mechanism for the impact of abnormal DNA methylation on transcriptional control resulting in cancers. We start from exploratory bioinformatics analyses of human genome non-coding regions, using next generation sequencing (NGS) data. We identify for the breast cancer cell line MCF7 (25), a hypermethylated region within the promoter of the NDUFA13 gene, 130 b.p. from the TSS (Figure S1). The NDUFA13 gene encodes a NADH dehydrogenase enzyme, a part of the electron transport chain in mitochondria that can function as a tumour suppressor (23). The hypermethylation of the NDUFA13 promoter leads to the downregulation of the gene, which increases cell proliferation and subsequently leads to the onset of breast cancer (24). Using the TFregulomeR tool (27), we continue with mapping of the transcription factor (TF) response elements near the hypermethylated cites that match with the ChIP-seq signals identified for the region. The analysis reveals two response elements (REs) for the E2F1-DP1 heterodimer (TTTCCCGC) and the BZIP homodimer CEBPB (TTACACTA). The identified abnormal methylation sites are located within the E2F1-DP1-RE, in the linker region between the two TF dimers, and in the 3’-flanking sites of the CEBPB-RE (Figure 1).

Next, we design the E2F1-DP1-CEBPB-DNA enhanceosome complex and proceed with all-atom molecular dynamics (MD) simulations. We perform protein-DNA docking with the HDOCK server (31), using the CEBPB dimer crystal structure (PDB ID: 1GU4) and a model structure of the E2F1-DP1 dimer derived through homology modelling in YASARA (28, 29), with the crystal structure of homologous E2F4-DP2 dimer (PDB ID: 1CF7 (30)) as the template. We dock each protein dimer individually to B-DNA composed of the TF-RE and two adjacent flanking nucleotides (NNRENN). This yields among top-ten highest ranked docking poses, the protein-DNA complexes resembling the expected binding modes, with RMSD of heavy atoms of 3.2 and 3.4 Å, for CEBPB- and E2F1-DP1-DNA complexes, respectively (Figure S3). Then, using USCF Chimera (32) we combine the two docked complexes through superposition on B-DNA that covers the two REs and six flanking b.p. on either 5’- and 3’-sides, 49 b.p. in total, to obtain the complete E2F1-DP1-CEBPB-DNA enhanceosome system.

We proceed with microsecond long all-atom MD simulations for DNA alone and in complex with E2F1-DP1-CEBPB-, CEBPB-, E2F1-DP1-factors for wild type (WT) and methylated (ME) systems, i.e., eight systems in total. Following the MD simulations, we compare the average structures of each WT-system with the corresponding ME-counterpart (Figure 2A). The differences are local, located within the binding region of the E2F1-DP1 dimer, where the CpG methylation sites contribute to a DNA bending towards the major groove. The differences between the WT- and ME-systems are more significant when the E2F1-DP1 dimer is bound to DNA (Figures 2A and S4), which is illustrated by the RMSD values of 5.6 Å for the E2F1-DP1-DNA systems and 4.5 Å for E2F1-DP1-CEBPB-DNA systems, respectively.

**Figure 2:**
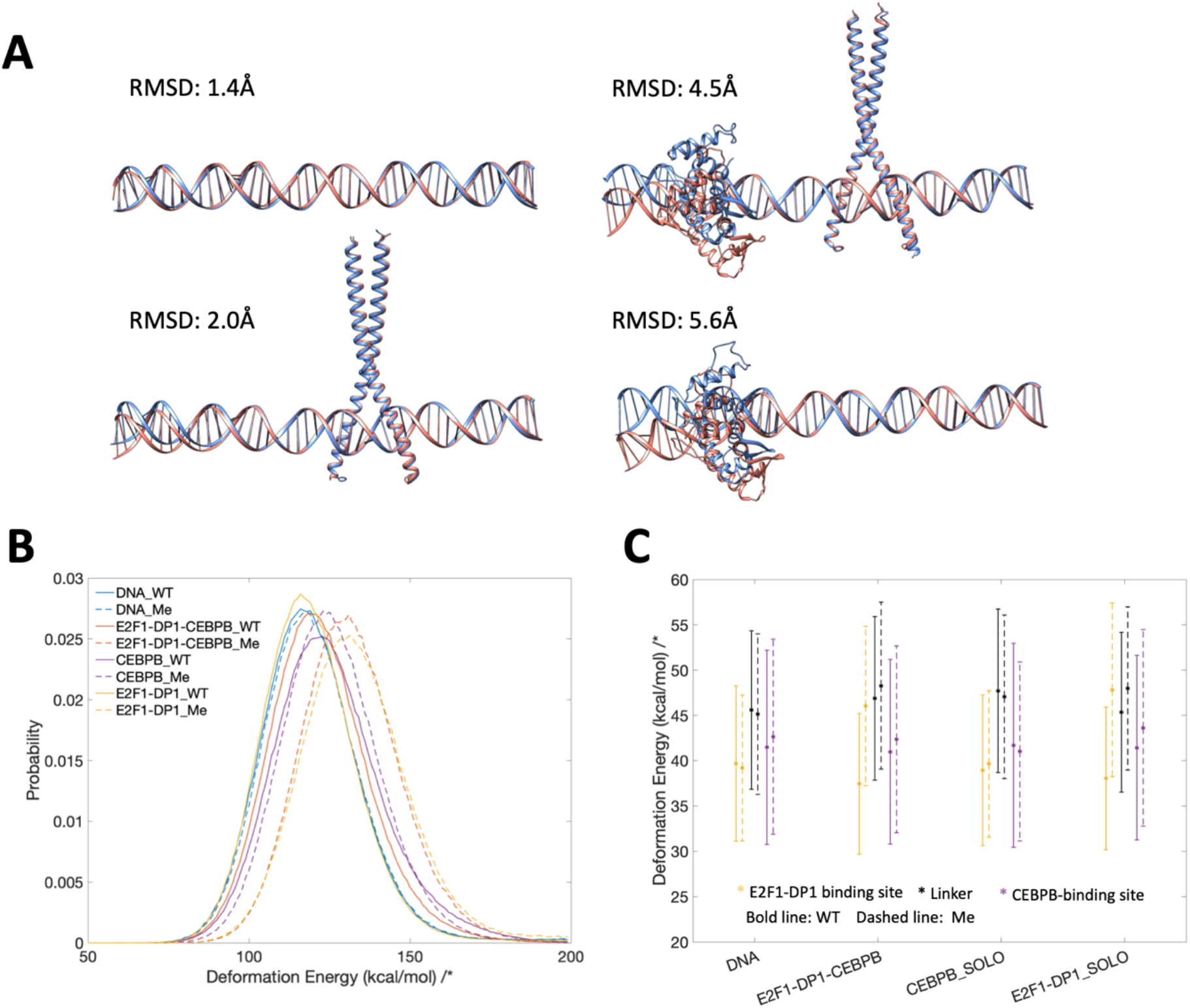
**A**. Comparison of average structures after 1 µs MD simulation for WT (blue) and Me (orange) systems. **B**. Distribution of DNA deformation energies for b.p. 6-35, which includes both response elements and the adjacent flanking nucleotides, calculated with a multivariate Isling model(43). **C**. DNA deformation energies for E2F1-DP1 binding site (b.p. 6-15), linker region (b.p.15-26), and CEBPB binding site (b.p.26-35).

### DNA deformation energies

Next, we analyze DNA deformation energy in all eight MD trajectories using a multivariate Ising model (43), derived to include the coupling between all possible sequence-specific conformational substates of DNA. Firstly, we calculate the deformation energy for the region covering both TF-REs and the first adjacent flanking nucleotides, e.g., 6-35 b.p. (Figure 2B, Table 1). We observe that in the WT-case, DNA deformation energy increases upon binding of the CEBPB-dimer to naked DNA, while the E2F1-DP1-DNA association has no obvious impact. However, when DNA becomes methylated, the E2F1-DP1-DNA binding further increases the deformation energy, irrespective if CEBPB is present or not. Secondly, we divide the DNA sequence into three regions: the E2F1-DP1 binding site, the linker, and the CEBPB binding site, and recalculate the deformation energy for each region. This segmentation provides further insights. In the WT-case, the CEBPB-DNA binding increases the deformation energy of the linker region, suggesting the induced conformational changes – analyses of DNA helical parameters confirm that (see below). Next, the E2F1-DP1-DNA binding increases the deformation energy of its DNA binding site and the linker region only in the ME-case. This suggests that cytosine methylation increases the energy cost of the E2F1-DP1-RE transition into its bioactive conformation. Taken together, the analyses hint at the cooperative nature of the TF-dimers-DNA association, which can be perturbed by the hypermethylation.

**Table 1:** DNA deformation energies (kcal/mol) for the 6-35 b.p. region, calculated using a multivariate Ising model (43).

|  | DNA | E2F1-DP1-CEBPB-DNA | CEBPB-DNA | E2F1-DP1-DNA |
| --- | --- | --- | --- | --- |
| **WT** | 120.0 ± 15.7 | 122.6 ± 15.7 | 125.5 ± 17.1 | 119.5 ± 15.6 |
| **ME** | 124.0 ± 15.8 | 132.3 ± 15.5 | 126.4 ± 15.4 | 134.5 ± 16.7 |

### Protein-DNA contacts

We continue with the analyses of protein-DNA contacts to further understand the mechanistic impact of CpG methylation on the E2F1-DP1-CEBPB-DNA enhanceosome formation. For the analyses we employ a dynamic contact map approach we derived earlier (41, 45), where for each MD frame we calculate a contact strength for every pair of protein-DNA residues that interact specifically and nonspecifically. We follow the time-evolution of the contacts strengths (Figures S5-S8) and calculate the average strength value for each protein-DNA contact (Figures 3, S9, S11, and S12). The analyses of the dynamic contact maps for the WT-vs ME-cases and solo-vs enhanceosome systems show that the CEBPB dimer exploits nearly the same network of contacts in all systems (Figures S9 and S11). This suggests that the CEBPB-DNA binding is independent of CpG methylation or the binding of the E2F1-DP1 dimer. The CEBPB dimer, is a BZIP factor, which recognizes its REs through specific interactions formed by a conserved five residues motif (NxxAVxxSR) (46), aar 281-289, of each monomer. The studied CEBPB-RE has a non-canonical CEBPB-half site (**TAGT**GTAA), which explains the observed differences in the specific contacts exploited by the two monomers (Figures S9 and S10). Of the five residues motif of monomer one, later CEBPB1, Ala284, Val285, and Ser288 interact hydrophobically with the flanking 5’-GpT, first TpA steps, and the adenine of the TpA step on the Crick strand (**GTA**GTGT**A**A), respectively, while Arg289 is involved in hydrogen bonding with the central CpA-step (TTA**CA**CTA) and the guanine of the first GpT step of the CEBPB1 half-site (TA**G**TGTAA). Though the Arg289 contacts are somewhat weaker in ME-systems. Specific interactions of the five residues motif of monomer two, later CEBP2, include cross-bridging hydrogen bonds by Asn281 to the first two TA b.p. (**T**TACACTA/TAGTGT**A**A), hydrophobic contacts by Ala284, Val285, and Ser288 with the flanking 5’-GpT step and the TpT step of the CEBPB2 half-site (**GTT**ACACTA), and hydrogen bonding by Arg289 with the central TpG step (TAG**TG**TAA). Some of the abovementioned contacts exploited by Ser288 and Arg289 of both CEBPB monomers only appear in some of the simulated systems (Figure S9-10). However, we believe that these contacts are not system specific, rather they are coupled to the flickering power of long-chained residues, which can fluctuate between different conformational substates.

**Figure 3.**
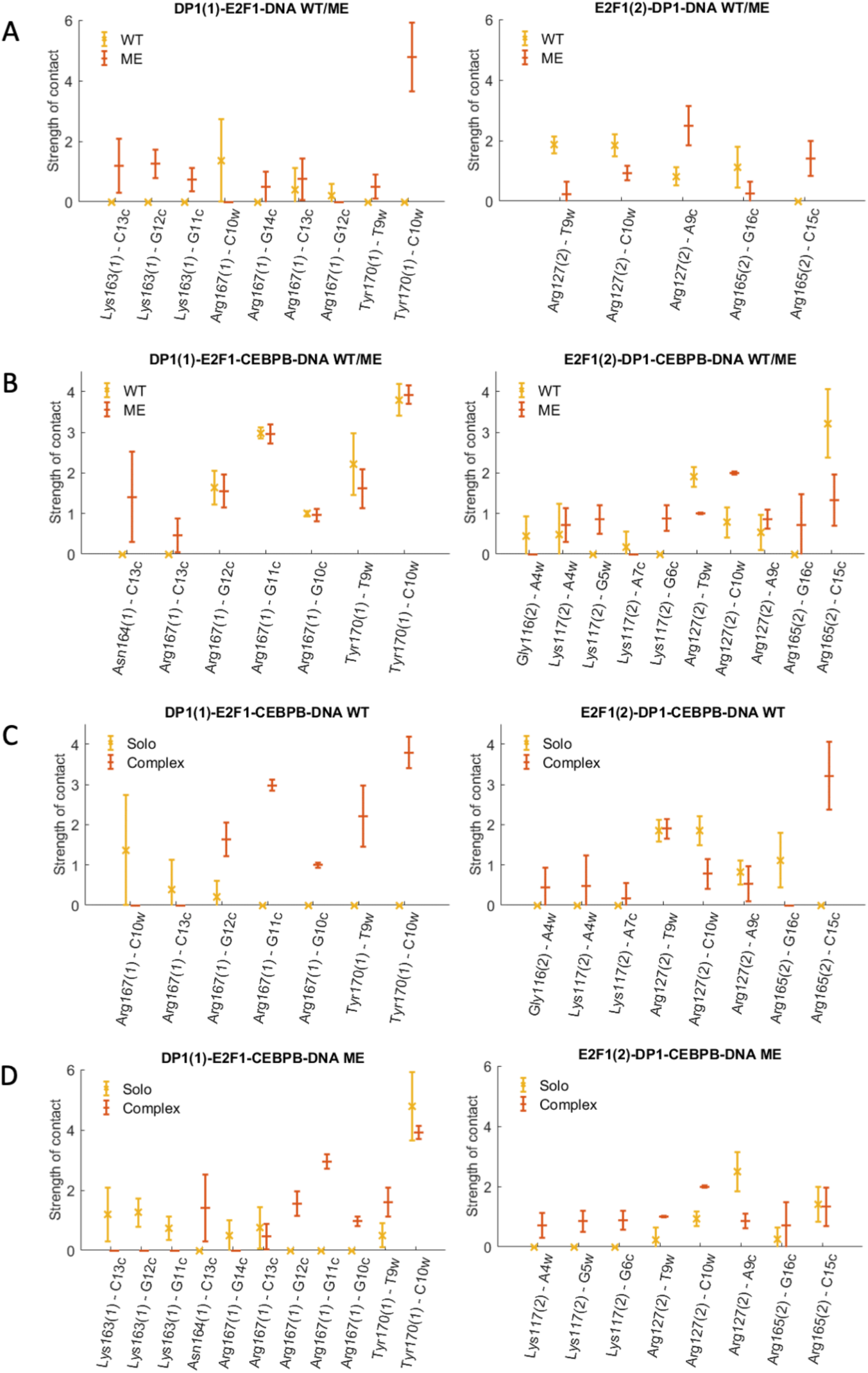
Specific contacts between E2F1-DP1 dimer and DNA. The plots show the strength of specific contacts formed by DP1(marked with (1)) and E2F1 (marked with (2)) monomers in wild-type (WT) and methylated (ME) systems, when bound alone to DNA (solo) and together with CEBPB (complex). For the definition of a contact strength see Supplementary Methods.

In contrast, the DNA-contacts exploited by the E2F1-DP1 dimer show a great variation among the simulated systems (Figures 3, 4A, and S12). We observe a rearrangement of the E2F1-DP1-DNA contacts when comparing WT- and ME-E2F1-DP1-DNA systems, and a strong impact of TF-cooperativity for the E2F1-DP1-DNA binding. The E2F1-DP1 dimer belongs to the E2F-DP family of TFs, the winged-helix folded proteins that recognize their REs with a conserved four residues motif (RRxYD), aar 167-171 and 165-169 for the DP1 and E2F1 monomers, respectively (30). For DP1, Arg167 is involved in hydrogen bonding with the three guanines of DNA Crick strand (TTTCCCGC/GC**GGG**AAA) in the WT-systems, in contrast the residue forms predominantly hydrophobic contacts with the methyl group of the C_W_13 base in ME-systems. Tyr170 forms hydrophobic contacts with the TpC step (TT**TC**CCGC) in WT-CEBPB-E2F1-DP1-DNA system and only with T_W_10 base (TT**T**CCCGC) in ME-E2F1-DP1-DNA system (Figure 4B). For E2F1, Arg165 forms hydrophobic contacts with the flanking GpC step (**GC**GCGGGAAA). For both monomers, Arg168/Arg166 interact nonspecifically with DNA backbone, while Asp171/Asp169 show no direct contacts with DNA, instead stabilizing the orientation of the Arg167-168/Arg165-166 residues. In addition, E2F1 has an N-terminal random coil tail that twists around DNA forming mostly nonspecific contacts, except for Arg127 which also specifically interacts with several b.p. of the RE (TT**TC**CCGC/CGGG**A**AA). Also, for WT- and ME-E2F1-DP1-CEBPB-DNA systems there is an additional specific contact by Lys117 of the N-terminal tail. Overall, we observe a reduction in the strength and the number of specific and nonspecific DNA-protein contacts for WT- and ME-E2F1-DP1-DNA systems (Figures 3 and S12). In WT-E2F1-DP1-DNA system, the DP1 recognition helix moves out of the major groove, while in the ME-case the recognition helix changes its orientation and slides downwards (Figure 4B). The two binding modes of the DP1 monomer in WT- and ME-E2F1-DP1-DNA systems lead to the loss of specificity of E2F1-DP1-DNA binding. In contrast, the presence of CEBPB secures a stable E2F1-DP1-DNA binding, with higher DNA sequence-specificity in the WT-case, which implies the importance of TF-cooperativity for the E2F1-DP1-CEBPB-DNA enhanceosome formation and suggests that the CEBPB-DNA binding should occur first.

**Figure 4.**
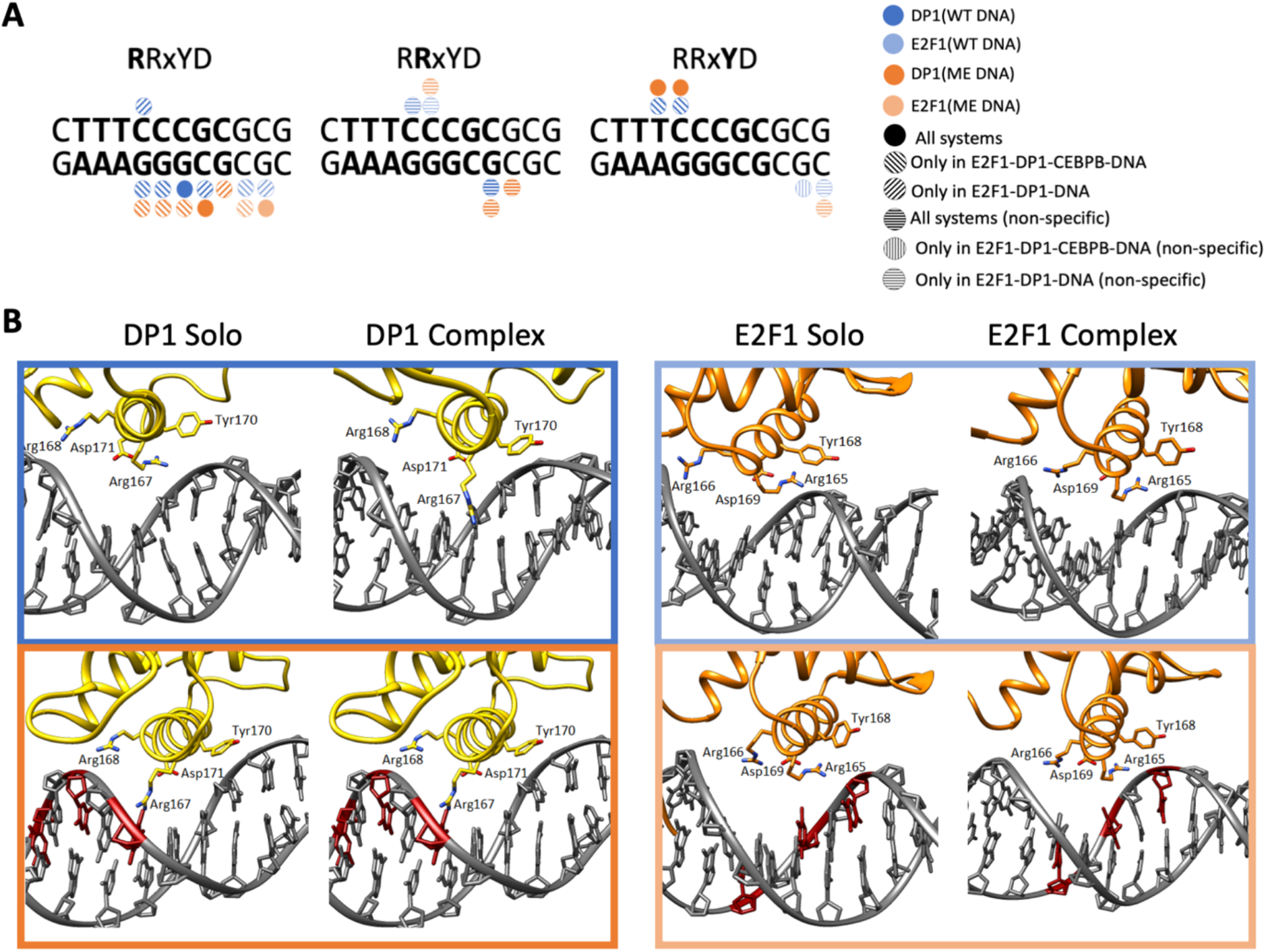
**A**: Schematic representation of specific protein-DNA contacts exploited by the four-residues-motif (RRxYD) of E2F1-DP1 in different systems. **B**: Cartoon representation of the binding orientations and DNA contacts of the E2F1-DP1 dimer for WT (blue) and ME (orange) systems, when bound alone to DNA (solo) and together with CEBPB (complex).

### Protein-DNA interaction energies

We proceed with the analysis of protein-DNA interaction energies. The accurate calculation of the protein-DNA interaction energy is difficult due to the massive negative charge of the DNA backbone. Nevertheless, for a qualitative comparison, we calculate the interaction energies including electrostatic and van der Waals, for specific and nonspecific protein-DNA contacts along all protein-bound trajectories using GROMACS energy analysis tool (Figures S13-S16). The interaction energies for the specific and nonspecific contacts follow the trends seen in the dynamic contact maps (Figures S5-S8). The interaction energy distributions for the specific contacts are bimodal (Figure S17), which can be coupled to the flickering behavior of the long side chain residues. The nonspecific interaction energy distributions (Figure S18) are normal and broad, which reflect the large variations in the number and contact strength of most of the nonspecific contacts. When we combine the interaction energies for specific and nonspecific contacts (Table 2), it appears that cytosine methylation contributes to a stronger TF-DNA complexation. However, the differences are small with overlapping standard deviations, which can be coupled to the different conformational substates exploited by the flickering residues. Instead, we see once again that the TF-cooperativity greatly affects the E2F1-DP1-DNA binding. We also calculate interaction-energies following the MMGBSA/MMPBSA approach (Table S1). We exclude calculations of conformational entropies due to the size of the systems. The MMGBSA/MMPBSA energies follow the same trends, again highlighting a cooperative dependency on CEBPB for strong E2F1-DP1-DNA interactions. The overlapping standard deviations make impossible any discussions on the effects of DNA methylation on the protein-DNA associations. Also, we should remember about the DNA deformation energies, which significantly increase when the methylation marks appear within the E2F1-DP1 binding site.

**Table 2.**
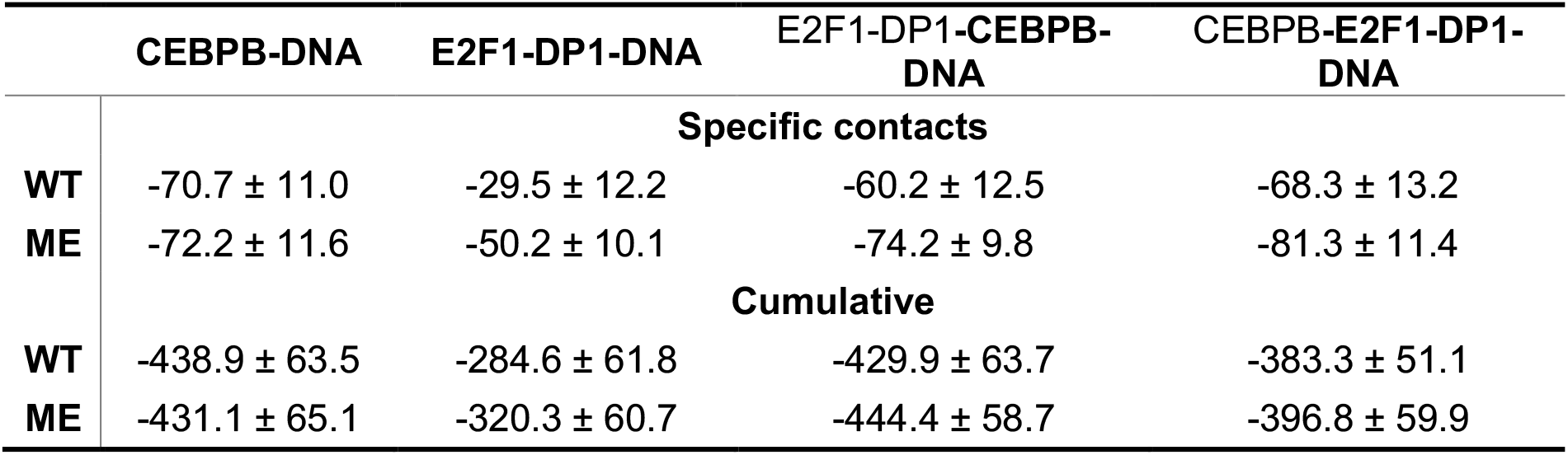
Protein-DNA interaction energies (kcal/mol) including standard deviations, calculated between all DNA-TF-dimer complexes. For E2F1-DP1-CEBPB-DNA trajectories, the provided interaction energies are for the protein dimer in bold.

### Changes in DNA helical and groove parameters

Next, we analyze changes in DNA helical and groove parameters for the WT-vs ME-cases and solo-vs enhanceosome systems (Figures 5 and S19-S25). Previously, we reported that the binding of proteins to their response elements results in small conformational adjustments in DNA helical parameters, which facilitates formation of stable specific contacts (41). The changes are reflected in reducing the bimodality/multimodality and narrowing of the helical parameters’ distributions, indicating that proteins stabilize a particular DNA substate. The analyses show that all CEBPB-bound DNA exhibit similar changes in helical and groove parameters, with respect to naked DNA, within the CEBPB-RE region (b.p. 27-34). This agrees with the protein-DNA contact analyses that show nearly identical CEBPB-DNA contact networks in all systems (Figure S10). In addition, the CEBPB binding introduces changes in shift, slide, and grooves parameters of the four flanking b.p. on each side of the CEBP-RE (b.p. 23-26 and 35-38), further in the linker region at C_W_20 and in the E2F1-DP1-RE at C_W_10 (Figure 5). The CpG20-21 step is not involved in any interactions with the E2F1-DP1 dimer but is a soft dinucleotide step conveniently located in the middle of the linker. The positive shift and the deeper major groove at CG10, in the presence of CEBPB, stabilize the hydrophobic contacts formed by Tyr170 of DP1, which secures the deep placement of the DP1 recognition helix into the major groove, thus explaining the nature of the observed TF cooperativity (Figure 4). In the ME-E2F1-DP1-CEBPB-DNA case, the effects from the CEBPB binding are smaller, due to the changes in helical shift coupled to DNA methylation (Figure 5).

**Figure 5.**
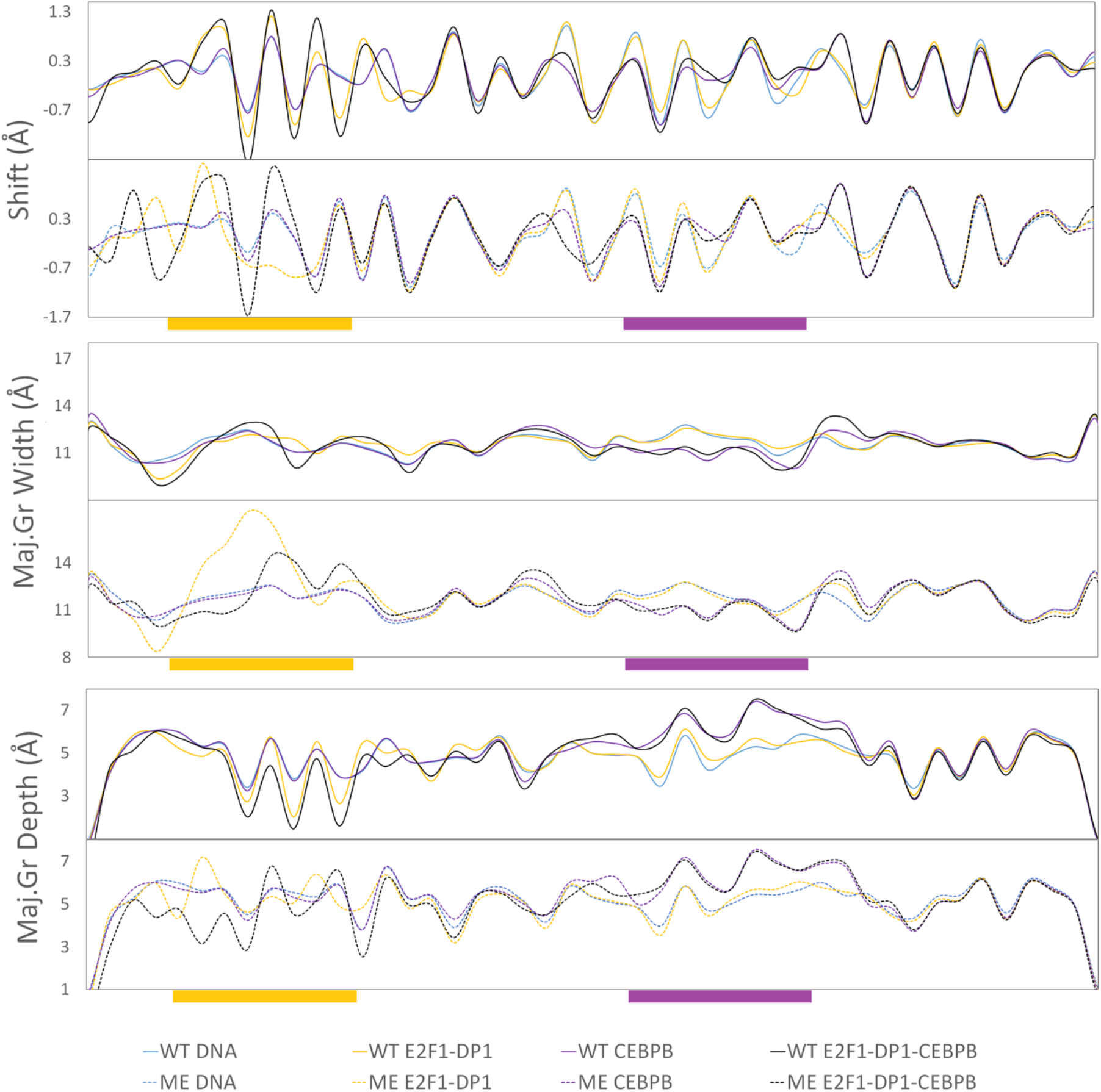
Average values for shift, major groove width and depth (in Å) for wild-type (WT) and methylated (ME) DNA alone and in complex with either CEBPB/E2F1-DP1 dimers, or complete CEBPB-E2F1-DP1. WT-values are depicted with bold lines, ME – with dashed lines.

In contrast, the changes in DNA helical and groove parameters induced upon the binding of the E2F1-DP1 dimer within its RE (7-14 b.p.), the flanking regions (b.p. 3-6 and 15-18), the linker, and further in the CEBPB-RE (b.p. 28, 30, 33-34) are system-specific. Similarly, to the CEBPB-DNA system, the changes induced by the E2F1-DP1 dimer happen at C_W_20. The changes in helical parameters within the CEBPB-RE, induced by the E2F1-DP1 binding, do not significantly alter the CEBPB-DNA specific contact network, as mentioned above. Methylation of CpG steps leads to a decreased twist and an increased roll (Figure S21-22), which through DNA backbone BI → BII transitions, induce changes in shift and slide, and consequently in major groove depth and width (Figures 5, S19-S20, S24-S25). Focusing on shift, where we observe the opposite sign of shift between the WT- and ME-cases for the E2F1-DP1-bound systems for the last three b.p. steps of E2F1-DP-RE (TTTCC**CGCG**). The effect is more obvious when we analyze the b.p. displacement in the x-axis (Figure S23). This explains the rearrangement of the DNA contacts formed by the E2F1-DP1 dimer, and the appearance of the new specific contacts by Lys163 and Asn164 of DP1 for the ME-E2F1-DP1-DNA and ME-E2F1-DP1-CEBPB-DNA systems, respectively (Figure 4). For the linker region, in addition to the change of the shift sign for the methylated b.p., we also observe the narrowing of the shift distributions (Figure S19 B). This indicates that DNA methylation stabilizes different from the WT-case conformational substates, which explains the damping of the allosteric communication between the two TF dimers.

Previous studies have shown that cooperativity between two TFs, which do not exhibit direct protein-protein interactions, arises due to DNA-mediated allosteric signals transmitted by one TF to the binding site of the other TF in a time-dependent manner (44). Thus, we next analyze long-distance correlations of DNA groove parameters and helical translational parameters with and without a time lag for the entire trajectories (0.1–1.1 µs) (Figures 6, S26, and S27). Our results show, in agreement with the previous studies, that the binding of CEBPB, but not E2F1-DP1, results in long-distance correlations of DNA b.ps. parameters (44). The strongest impact of the CEBPB binding on long distance correlations is seen for the shift parameter and the major groove width (Figure 6). The signal is translated from C_W_32 in the middle of the CEBPB1-RE, over to G_W_21G_W_22A_W_23 and then from T_W_19C_W_20 in the linker region to the E2F1-DP1-RE b.p. 10-14 (TTT**CCCGC**). However, the correlation coefficients are small (< 0.3), therefore caution should be taken. The addition of a time lag has not improved the long-distance correlations. Furthermore, the long-distance communication between the two binding sites increases when both TFs are bound to DNA. This could indicate that cooperativity and communication firstly arise when one TF is bound to DNA and the other slides and approaches its binding site. The long-distance correlations somewhat increase (< 0.35) when calculated for the trajectories intervals when the flickering residues form specific contacts with DNA. However, we believe that taking the complete trajectories provides a more complete picture of the process. The described long-distance correlations observed within the linker and E2F1-DP1-RE regions upon the CEBPB binding are missing when DNA becomes methylated (Figures S26-S27), suggesting that methylation hinders the propagation of the allosteric signal.

**Figure 6.**
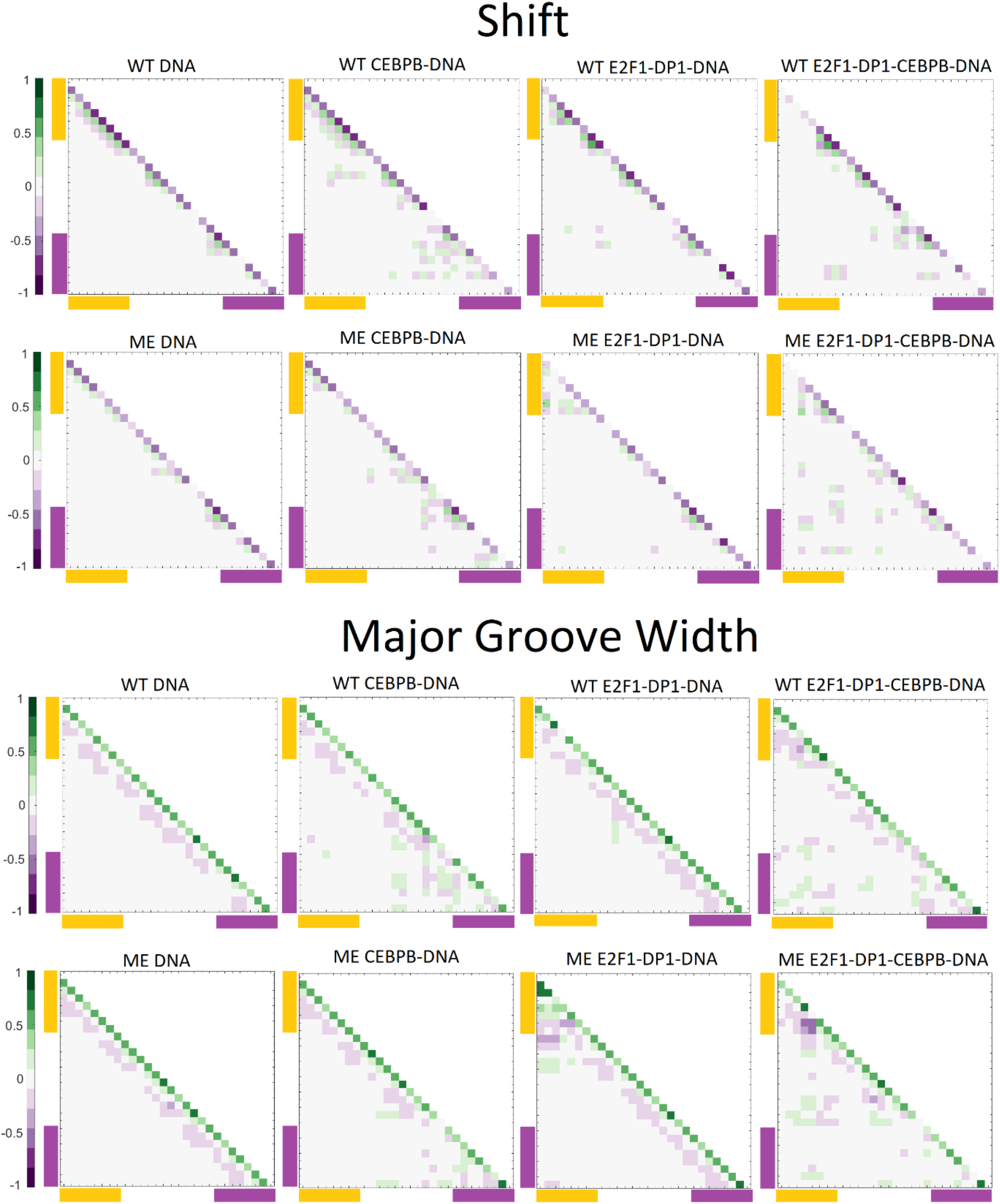
Correlation coefficient maps without time lag for b.p. shift and major groove width along the WT and ME sequences for DNA alone and in complex with either CEBPB/E2F1-DP1 dimers, or with both TFs dimers CEBPB-E2F1-DP1. Correlation coefficients are calculated for the entire (0.1–1.1 µs) trajectories. In all panels, the E2F1-DP1 response element is marked with yellow, the CEBPB response element – with magenta.

### Ion Populations

Previous studies have shown that counterion populations within DNA grooves are sequence-specific: depending both on sequence-specific flexibility of DNA backbone and electronegative groups in DNA bases, and may impact the TF-DNA association (47–49). Thus, we also investigate the counterion populations within the major and minor grooves for naked and protein-bound WT- and ME-DNA systems, using Canion program. Firstly, for WT-systems, we observe an elevated K+ concentration at C10 and C14 positions in the absence of CEBPB (Figure S28 A). Secondly, we observe a slight increase in K+ molarity in the major groove of the E2F1-DP1-RE region for naked and CEBPB-bound ME-DNA, in comparison to the corresponding WT-systems. Additionally, the analyses show that cytosine methylation affects the radial distribution of K+ ions within the major groove; the ions are shifted 0.5-1.0 Å closer to DNA helical axis in respect to WT-DNA (Figure S28 B). These observations agree with the conclusions derived through dynamic contact maps showing that the recognition helix of DP1 moves out of the major groove in the WT-E1F2-DP1-DNA case, and in the ME-E1F2-DP1-DNA case the helix slides downwards reducing the specificity of the DP1-DNA binding.

## Discussion

Using NGS data on breast cancer cell line MCF7, we have investigated the mechanistic impact of abnormal DNA methylation in the non-coding genome on the downregulation of NDUFA13 gene. The downregulation of NDUFA13 gene, which encodes a tumor suppressor NADH dehydrogenase enzyme, is observed in multiple cancers (11, 23, 24). We have identified a hypermethylated DNA region 130 b.p. from the NDUFA13 gene TSS, which also contains the DNA binding sites for two TFs dimers, CEBPB and E2F1-DP1. The proximity to the TSS and overlap with the TFs-REs suggest that this hypermethylated site may contribute to the downregulation of NDUFA13 gene. To verify the hypothesis, we have performed all-atom microsecond long MD simulation of wild-type and methylated DNA alone and in complex with either solo CEPBP/E2F1-DP1 dimers, or E2F1-DP1-CEBPB-DNA enhanceosome systems. We analyzed protein-DNA interactions, changes in DNA conformational dynamics, and ion distributions. Our data allow to describe with atom-level detail the mechanism of cooperative DNA recognition by the two TF dimers, and how the process is affected by DNA hypermethylation.

In terms of cooperative DNA recognition, we see that the primary binding of CEBPB stabilizes the E2F1-DP1-DNA association. This is reflected by the stronger E2F1-DP1-DNA interaction energies and by the binding orientation of the E2F1-DP1 dimer, deeper within DNA major groove, in the presence of CEBPB. Through alterations in DNA helical and groove parameters in the linker region, the primary CEBPB-DNA binding induces local structural changes within the E2F1-DP1-RE that facilitate the formation of the specific contacts by Tyr170 of DP1. Our data suggest that Tyr170-DNA specific contacts anchor the E2F1-DP1 dimer into DNA major groove. This agrees well with a recent published statistical analysis of SELEX data that identifies DNA shape as a major component of TF-cooperativity (50).

Interestingly, the primary E2F1-DP1-DNA binding also induces changes in DNA helical and groove parameters within the CEBPB-RE, however they do not noticeably affect the CEBPB-DNA contact network. This agrees with our previous study that shows that BZIP TFs possess a great adaptability to changes in DNA structure (45). This in turn, allows us to hypothesize that CEBPB-DNA binding will happen first, which will facilitate the E2F1-DP1-CEBPB-DNA enhanceosome formation. However, the strongest DNA-mediated communication between the TFs dimers, through increased number of long-distance correlations of helical and groove parameters, arises only when both TFs dimers are bound to DNA. This is different from the previous published mechanism for the BAMHI-GRDBD cooperativity (44), which arises due to time-dependent allosteric signals transmitted by BAMHI to the binding site of GRDBD. This illustrates that there is no universal mechanism of TF-cooperativity. In the case of E2F1-DP1-CEBPB cooperativity, the DNA-mediated allosteric communication between two binding sites might first be senesced when CEBPB is bound to DNA and E2D1-DP1 approaching its response element.

In terms of the CpG methylation impact, E2F1-DP1 has been reported to be deficient in binding to methylated DNA (51). However, the ability of E2F1-DP1 to bind DNA may depend on DNA sequence-specific flexibility and whether the binding site contains a single or a double CpG methylation. The here-studied DNA sequence contain single methylated CpG marks within the E2F1-DP1-RE and linker regions, and a double methylated CpG in the 3’-flanks of the CEBPB-RE. Our data show that CpG methylation within the E2F1-DP1-RE leads to different conformational substates in DNA helical and groove parameters, making DNA wider and shallower in the major groove, which results in the loss of specificity for some E2F1-DP1-DNA contacts, but overall, a stronger E2F1-DP1-DNA association. Yet, our analyses reveal that the DNA deformation energy increases significantly in the ME-E2F1-DP1-DNA system, in comparison with the WT-case. This suggests that the E2F1-DP1 association with methylated DNA will cost more energy, which may imply that the E2F1-DP1 will no longer recognize it as a true binding site (52). Furthermore, CpG methylation affects the structural changes within the E2F1-DP1-RE induced by the CEBPB-DNA binding, which is reflected, e.g., by the loss of bimodality in shift and groove parameters in the linker region. We also observe that the long-distance correlations of DNA helical and groove parameters, seen for the WT-case, disappear. Finally, the double CpG methylation in the 3’-flanks of the CEBPB-RE seem to have no effect on either the CEBPB- or E2F1-DP1-DNA binding. Taken together, our data suggest that abnormal methylation in non-coding genome may impede the cooperative DNA recognition by collaborative TF proteins, which will impact gene transcription. We thus propose that the identified hypermethylated region in the promoter of NDUFA13 gene can be used as a biomarker (22) for breast cancer.

## Data availability

All data generated and analyzed in this study are available from the corresponding author upon request.

## Supplementary data

Supplementary Data are available at NAR online.

## Acknowledgements

The authors thank Swedish National Infrastructure for Computing (SNIC) for the generous provision of computing resources.

