## Supplementary material for "Methylation in NDUFA13 gene promoter disrupts communication between collaborative transcription factors – potential mechanism for onset of breast cancer"

### Supplementary methods

#### Protein-DNA contacts network analysis

Analysis of protein-DNA contacts for the E2F1-DP1-DNA/CEBPB-DNA/E2F1-DP1-CEBPB-DNA complexes is performed using CPPTRAJ16, for each trajectory snapshot extracted at 1 ps intervals. Protein-DNA contacts present for less than 10% of the trajectories are excluded. Protein-DNA contacts are characterized by pairs of residues, divided into 'specific', i.e., interactions formed between the protein side chains and DNA bases, and 'nonspecific', i.e., interactions formed with at least one of the backbones. The contacts formed between each protein-DNA residue pair are summed, where for simplicity, the contribution of each contact is set to 1. The distance limit of a hydrogen bond interaction is ≤ 4 Å between the relevant heavy atoms, and the angle limit is ≥ 135° at the intervening hydrogen atom. For a hydrophobic contact, the limit is ≤ 6Å between the centre of mass of hydrophobic residues (Ala, Ile, Leu, Met, Phe, Trp, and Cys) and DNA bases. The derived time series of protein-DNA interactions allow construction of dynamic contacts maps for specific and nonspecific contacts.

### Supplementary figures

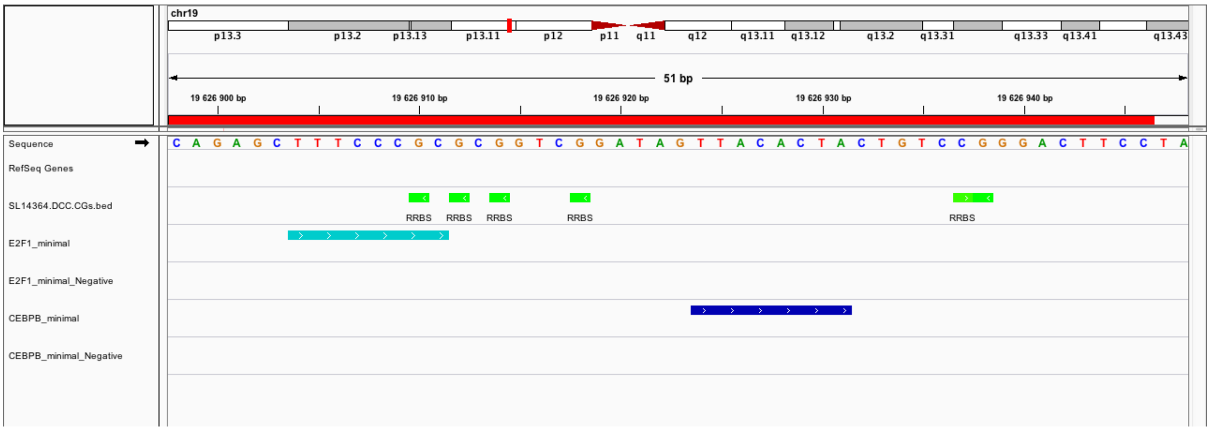


**Figure S1.** IGV (Integrative Genome Viewer) screenshot depicting the studied wild-type DNA sequence with the cytosine methylation marks detected in the promoter of NDUFA13 gene of breast cancer cell line MCF7; binding sites of two collaborative transcription factor dimers E2F1-DP1 and CEBPB.

**A**

```
E2F1          121 SPGEKSRYETSLNLTTKRFLELLSHSADGVVDLNWAAEVLKV-QKRRIYD
                  .||..||:|.||.|.|.:|:.||..:.|||:||..||:.|.| |||||||
E2F4           11 PPGTPSRHEKSLGLLTTKFVSLLQEAKDGVLDLKLAADTLAVRQKRRIYD

E2F1              ITNVLEGIQLIAKKSKNHIQWLGSH 194
                  ||||||||.||.|||||.|||.|..
E2F4              ITNVLEGIGLIEKKSKNSIQWKGVG 85
```

**B**

```
DP1           113 GKGLRHFSMKVCEKVQRKGTTSYNEVADELVAEFSAADNHILPNESAYDQ
                  |||||||||||||||||||||||||||||||:||:.::|| |..:|||||
DP2           129 GKGLRHFSMKVCEKVQRKGTTSYNEVADELVSEFTNSNNH-LAADSAYDQ

DP1               KNIRRRVYDALNVLMAMNIISKEKKEIKWIGLP 195
                  |||||||||||||||||||||||||||||||||
DP2               KNIRRRVYDALNVLMAMNIISKEKKEIKWIGLP 210
```

**Figure S2:** Pairwise sequence alignment used for homology modelling of the E2F1-DP1 heterodimer. **A.** Alignment between E2F1 (Query) and E2F4 (Template). **B.** Alignment between DP1 (Query) and DP2 (Template).

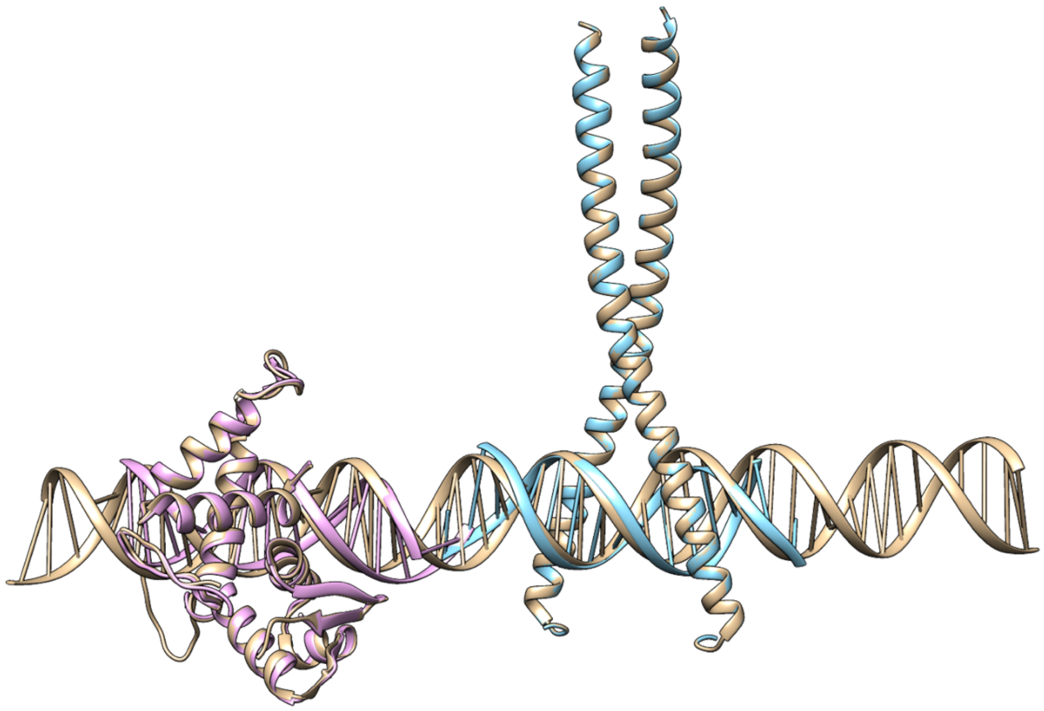

**Figure S3:** E2F4-DP2 (pink) and 1GU4 (cyan) PDB crystal structures superposed on the complete E2F1-DP1-CEBPB molecular model after homology modelling and docking to the DNA.

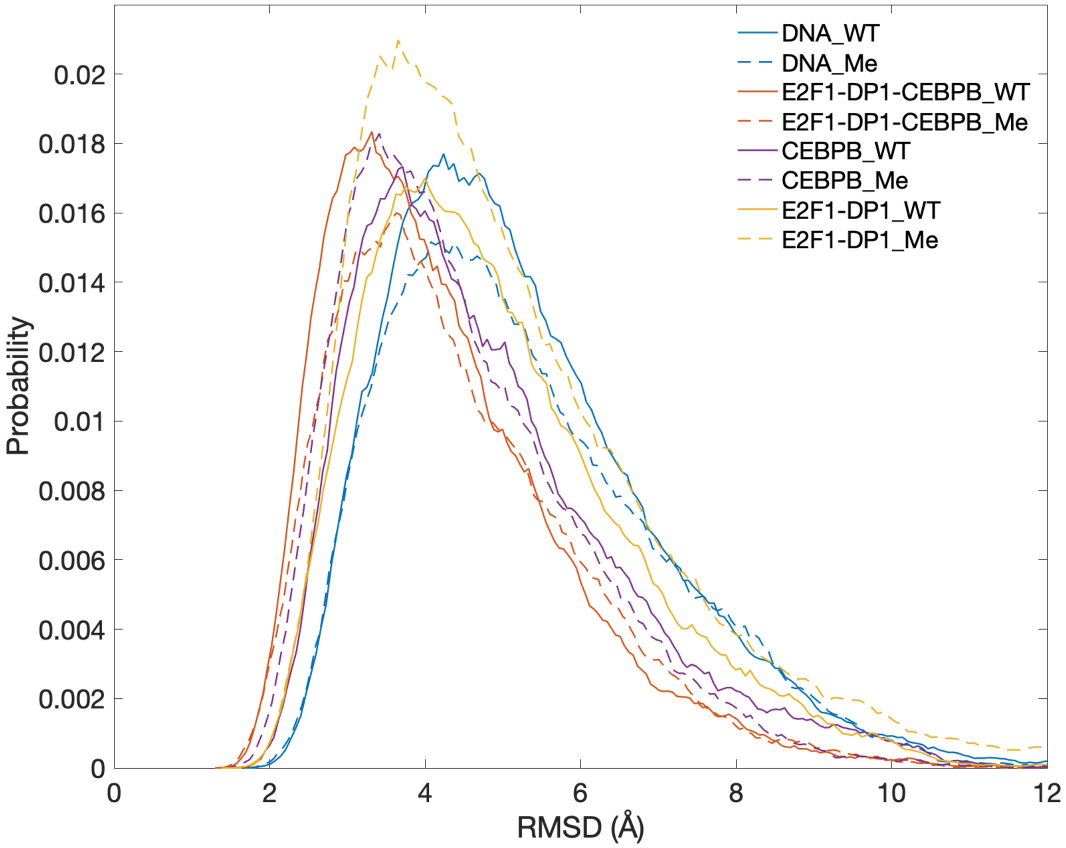


**Figure S4:** RMSD Distributions for the heavy atoms of DNA in different simulated systems.

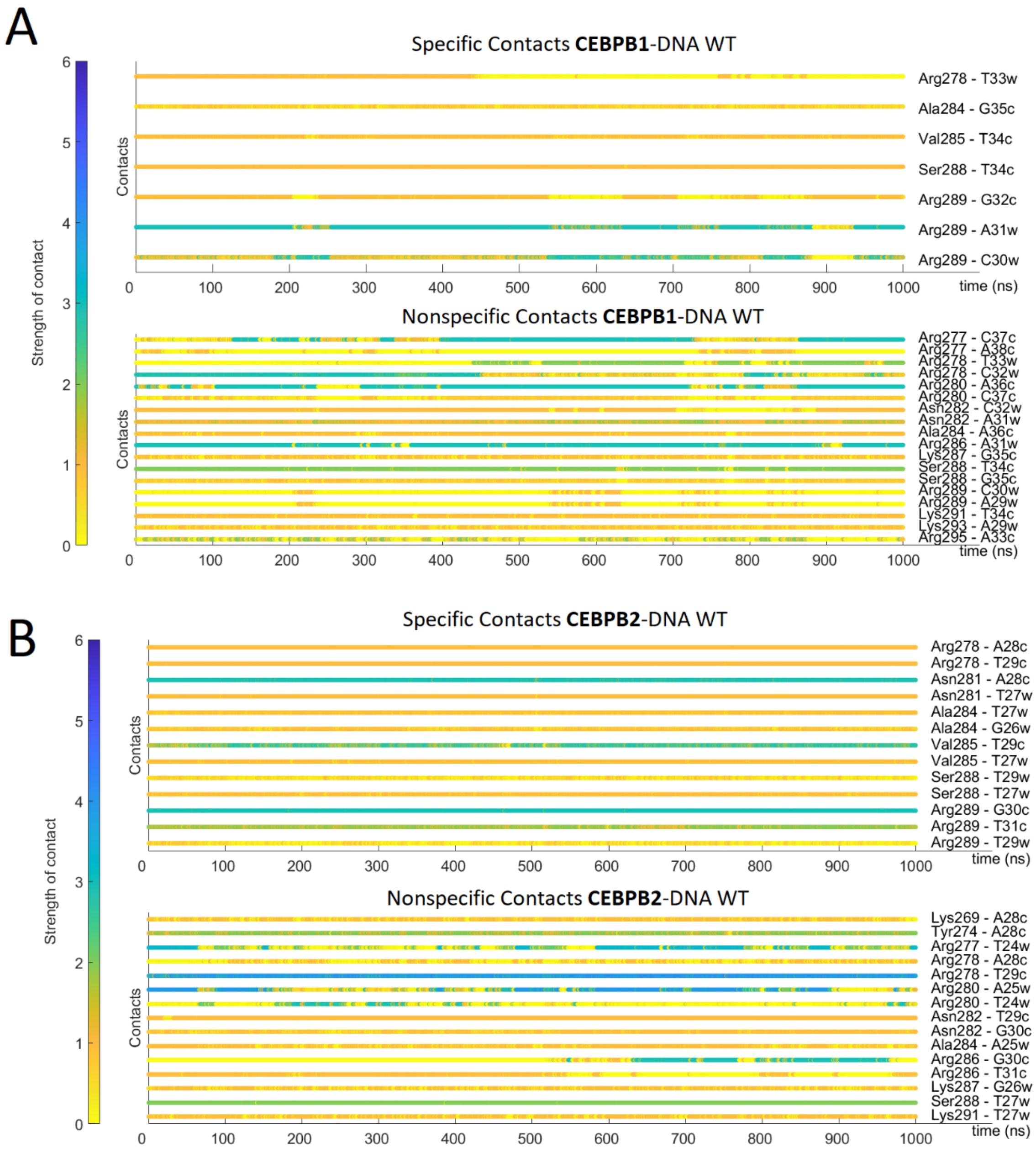
A
Specific Contacts CEBPB1-DNA WT
Strength of contact
Contacts
time (ns)
Arg278 - T33w
Ala284 - G35c
Val285 - T34c
Ser288 - T34c
Arg289 - G32c
Arg289 - A31w
Arg289 - C30w
Nonspecific Contacts CEBPB1-DNA WT
Arg277 - C37c
Arg277 - A38c
Arg278 - T33w
Arg278 - C32w
Arg280 - A36c
Arg280 - C37c
Asn282 - C32w
Asn282 - A31w
Ala284 - A36c
Arg286 - A31w
Lys287 - G35c
Ser288 - T34c
Ser288 - G35c
Arg289 - C30w
Arg289 - A29w
Lys291 - T34c
Lys293 - A29w
Arg295 - A33c
B
Specific Contacts CEBPB2-DNA WT
Arg278 - A28c
Arg278 - T29c
Asn281 - A28c
Asn281 - T27w
Ala284 - T27w
Ala284 - G26w
Val285 - T29c
Val285 - T27w
Ser288 - T29w
Ser288 - T27w
Arg289 - G30c
Arg289 - T31c
Arg289 - T29w
Nonspecific Contacts CEBPB2-DNA WT
Lys269 - A28c
Tyr274 - A28c
Arg277 - T24w
Arg278 - A28c
Arg278 - T29c
Arg280 - A25w
Arg280 - T24w
Asn282 - T29c
Asn282 - G30c
Ala284 - A25w
Arg286 - G30c
Arg286 - T31c
Lys287 - G26w
Ser288 - T27w
Lys291 - T27w

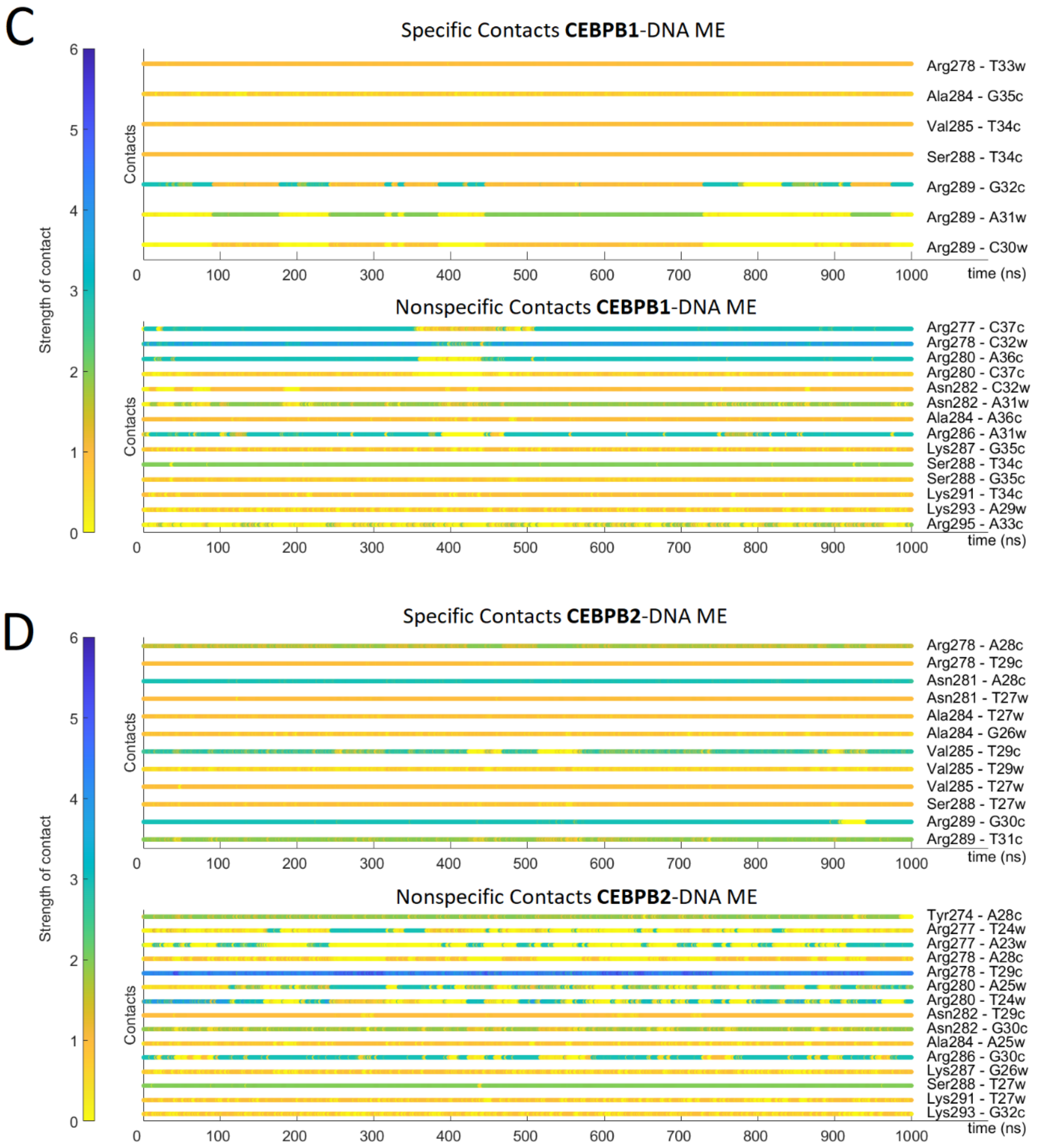


**Figure S5.** Time-evolution of the specific and nonspecific protein-DNA contacts for the CEBPB-DNA system, CEBPB1 and CEBPB2 correspond to different monomers: **A:** WT CEBPB1-DNA; **B:** WT CEBPB2-DNA; **C:** ME CEBPB1-DNA; **D:** ME CEBPB2-DNA.

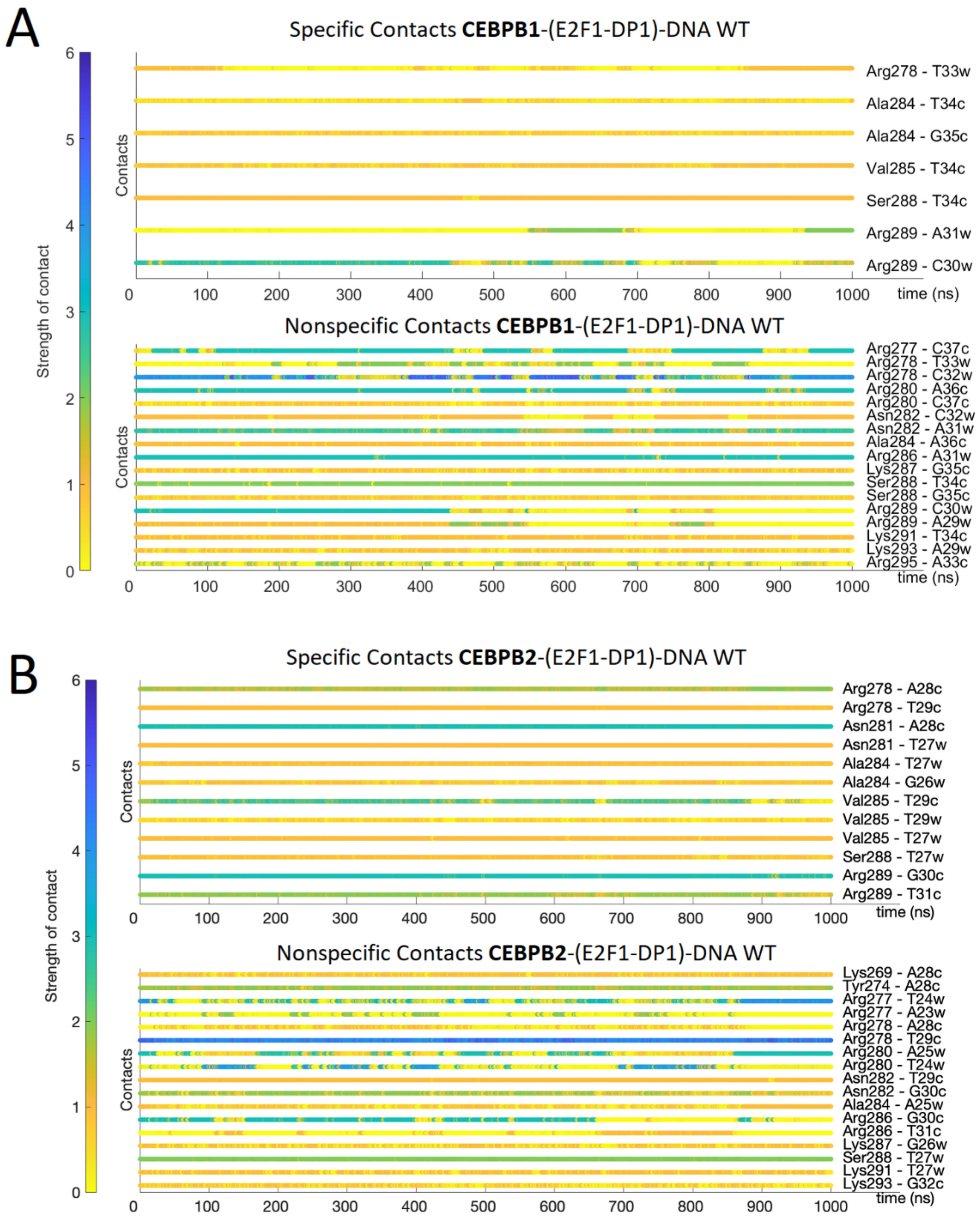
A
Specific Contacts CEBPB1-(E2F1-DP1)-DNA WT
Strength of contact
Contacts
Arg278 - T33w
Ala284 - T34c
Ala284 - G35c
Val285 - T34c
Ser288 - T34c
Arg289 - A31w
Arg289 - C30w
time (ns)
Nonspecific Contacts CEBPB1-(E2F1-DP1)-DNA WT
Arg277 - C37c
Arg278 - T33w
Arg278 - C32w
Arg280 - A36c
Arg280 - C37c
Asn282 - C32w
Asn282 - A31w
Ala284 - A36c
Arg286 - A31w
Lys287 - G35c
Ser288 - T34c
Ser288 - G35c
Arg289 - C30w
Arg289 - A29w
Lys291 - T34c
Lys293 - A29w
Arg295 - A33c
0 100 200 300 400 500 600 700 800 900 1000
B
Specific Contacts CEBPB2-(E2F1-DP1)-DNA WT
Arg278 - A28c
Arg278 - T29c
Asn281 - A28c
Asn281 - T27w
Ala284 - T27w
Ala284 - G26w
Val285 - T29c
Val285 - T29w
Val285 - T27w
Ser288 - T27w
Arg289 - G30c
Arg289 - T31c
Nonspecific Contacts CEBPB2-(E2F1-DP1)-DNA WT
Lys269 - A28c
Tyr274 - A28c
Arg277 - T24w
Arg277 - A23w
Arg278 - A28c
Arg278 - T29c
Arg280 - A25w
Arg280 - T24w
Asn282 - T29c
Asn282 - G30c
Ala284 - A25w
Arg286 - G30c
Arg286 - T31c
Lys287 - G26w
Ser288 - T27w
Lys291 - T27w
Lys293 - G32c

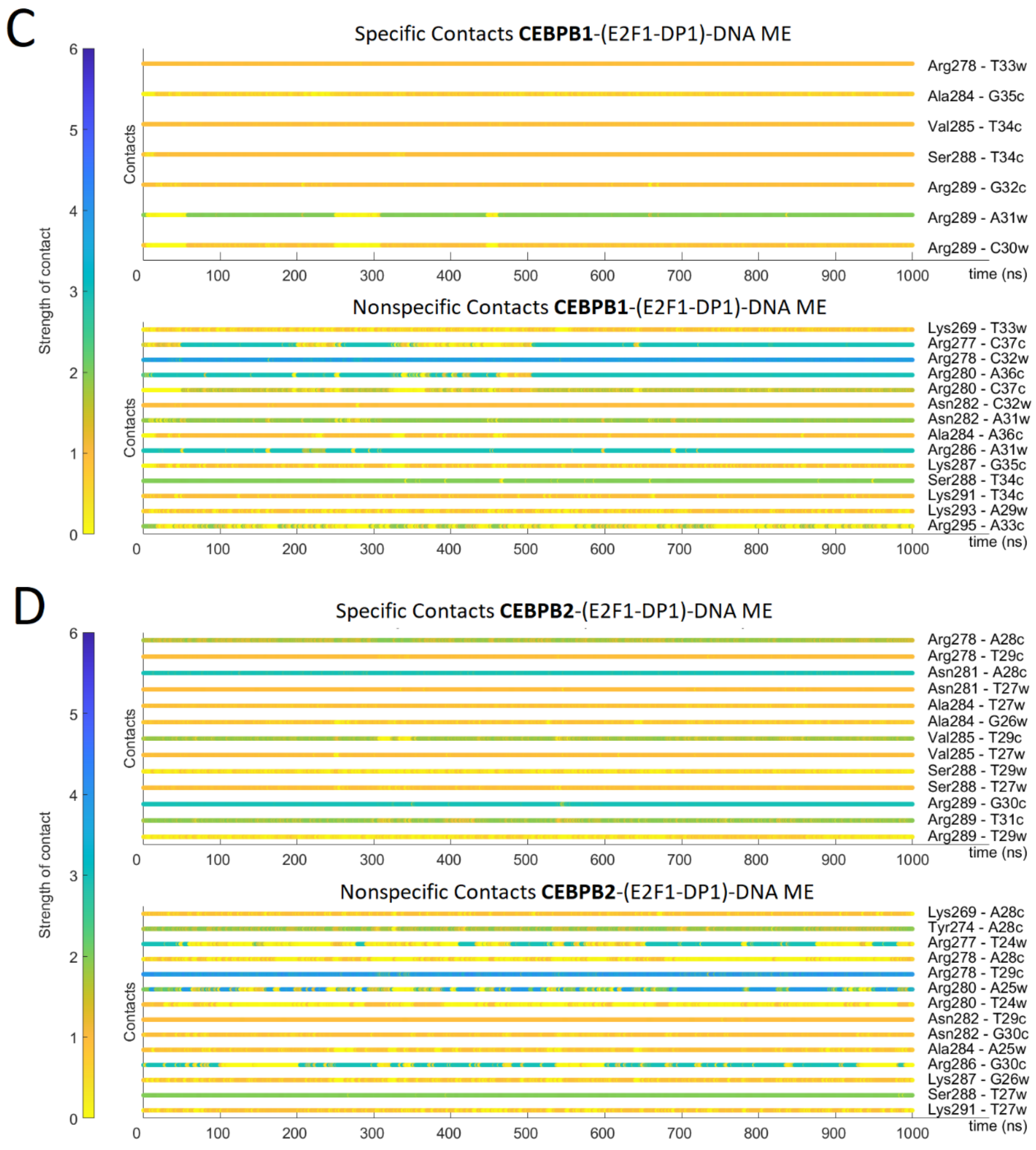


**Figure S6.** Time-evolution of the specific and nonspecific protein-DNA contacts for the E2F1-DP1-CEBPB-DNA system: **A:** WT CEBPB1-DNA; **B:** WT CEBPB2-DNA; **C:** ME CEBPB1-DNA; **D:** ME CEBPB2-DNA.

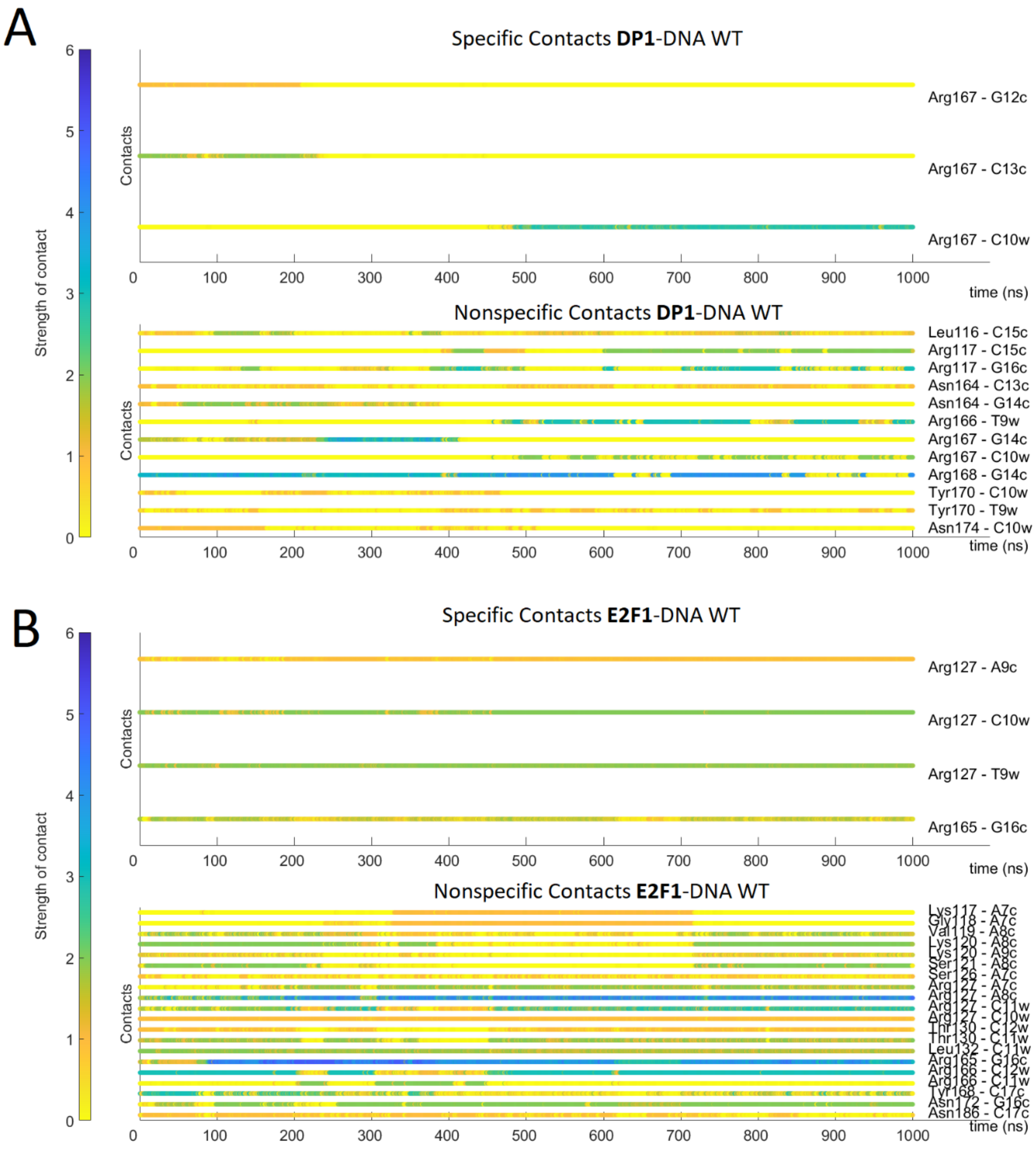
A
Specific Contacts DP1-DNA WT
Arg167 - G12c
Arg167 - C13c
Arg167 - C10w
Nonspecific Contacts DP1-DNA WT
Leu116 - C15c
Arg117 - C15c
Arg117 - G16c
Asn164 - C13c
Asn164 - G14c
Arg166 - T9w
Arg167 - G14c
Arg167 - C10w
Arg168 - G14c
Tyr170 - C10w
Tyr170 - T9w
Asn174 - C10w
Contacts
time (ns)
Strength of contact
B
Specific Contacts E2F1-DNA WT
Arg127 - A9c
Arg127 - C10w
Arg127 - T9w
Arg165 - G16c
Nonspecific Contacts E2F1-DNA WT
Lys117 - A7c
Gly118 - A7c
Val119 - A8c
Lys120 - A8c
Lys120 - A9c
Ser121 - A8c
Ser126 - A7c
Arg127 - A7c
Arg127 - A8c
Arg127 - C11w
Arg127 - C10w
Thr130 - C12w
Thr130 - C11w
Leu132 - C11w
Arg165 - G16c
Arg166 - C12w
Arg166 - C11w
Tyr168 - C17c
Asn172 - G16c
Asn186 - C17c
Contacts
time (ns)
Strength of contact

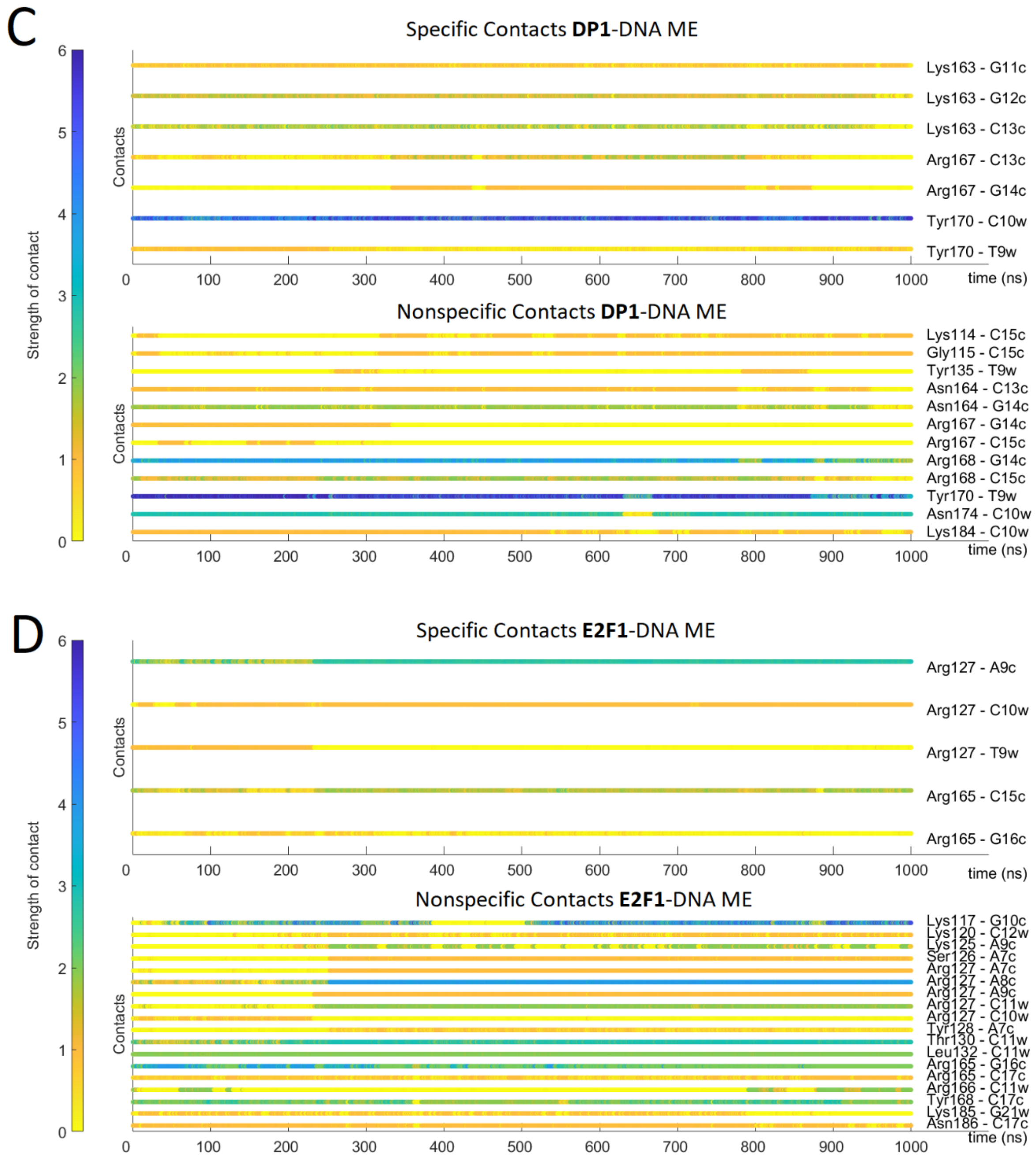


**Figure S7.** Time-evolution of the specific and nonspecific protein-DNA contacts for the E2F1-DP1-DNA system: **A:** WT DP1-DNA; **B:** WT E2F1-DNA; **C:** ME DP1-DNA; **D.** ME E2F1-DNA.

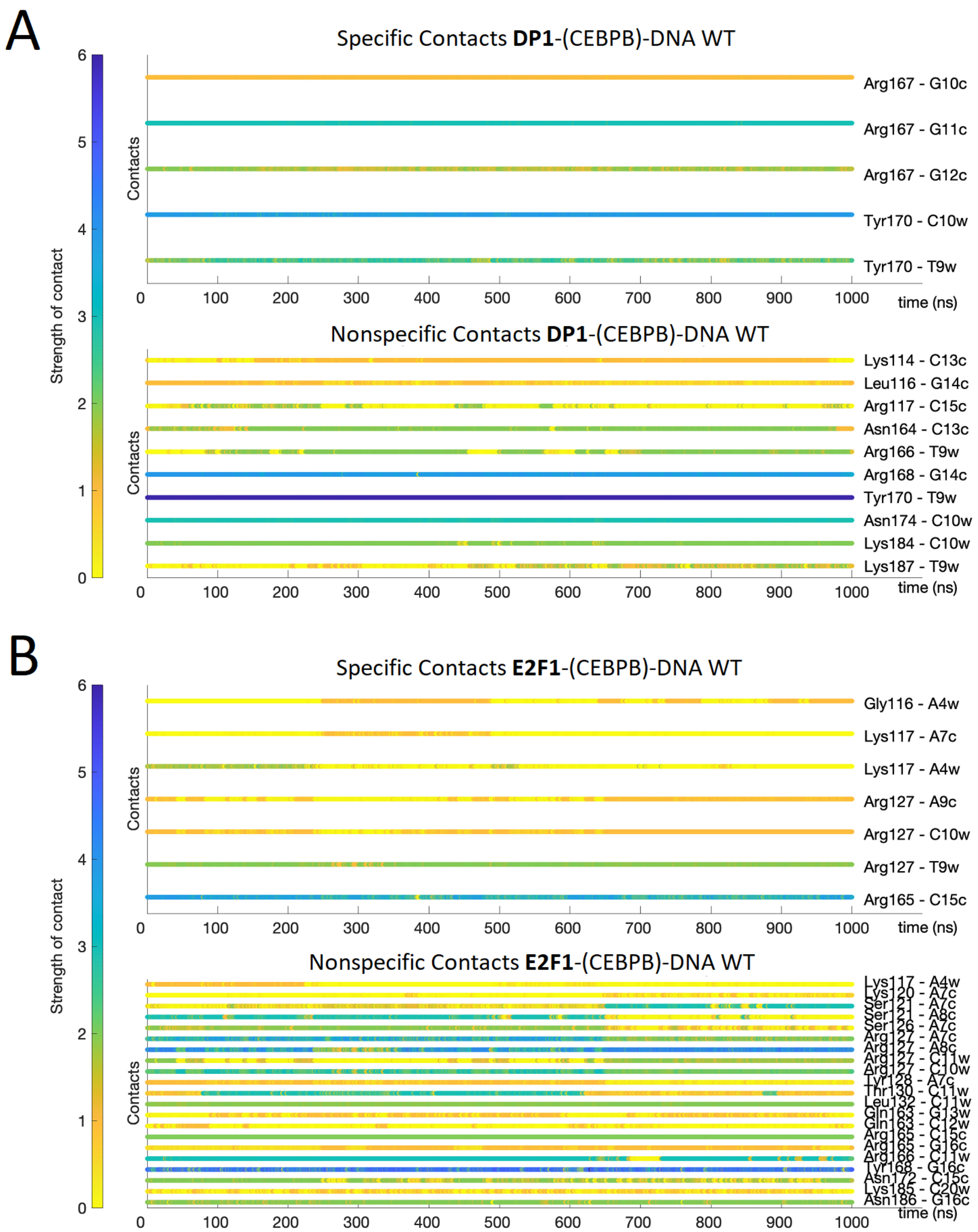
A
Specific Contacts DP1-(CEBPB)-DNA WT
Strength of contact
Contacts
Arg167 - G10c
Arg167 - G11c
Arg167 - G12c
Tyr170 - C10w
Tyr170 - T9w
time (ns)
Nonspecific Contacts DP1-(CEBPB)-DNA WT
Lys114 - C13c
Leu116 - G14c
Arg117 - C15c
Asn164 - C13c
Arg166 - T9w
Arg168 - G14c
Tyr170 - T9w
Asn174 - C10w
Lys184 - C10w
Lys187 - T9w
B
Specific Contacts E2F1-(CEBPB)-DNA WT
Gly116 - A4w
Lys117 - A7c
Lys117 - A4w
Arg127 - A9c
Arg127 - C10w
Arg127 - T9w
Arg165 - C15c
Nonspecific Contacts E2F1-(CEBPB)-DNA WT
Lys117 - A4w
Lys120 - A7c
Ser121 - A7c
Ser121 - A8c
Ser126 - A7c
Arg127 - A7c
Arg127 - A8c
Arg127 - C11w
Arg127 - C10w
Tyr128 - A7c
Thr130 - C11w
Leu132 - C11w
Gln163 - G13w
Gln163 - C12w
Arg165 - C15c
Arg165 - G16c
Arg166 - C11w
Tyr168 - G16c
Asn172 - C15c
Lys185 - C20w
Asn186 - G16c

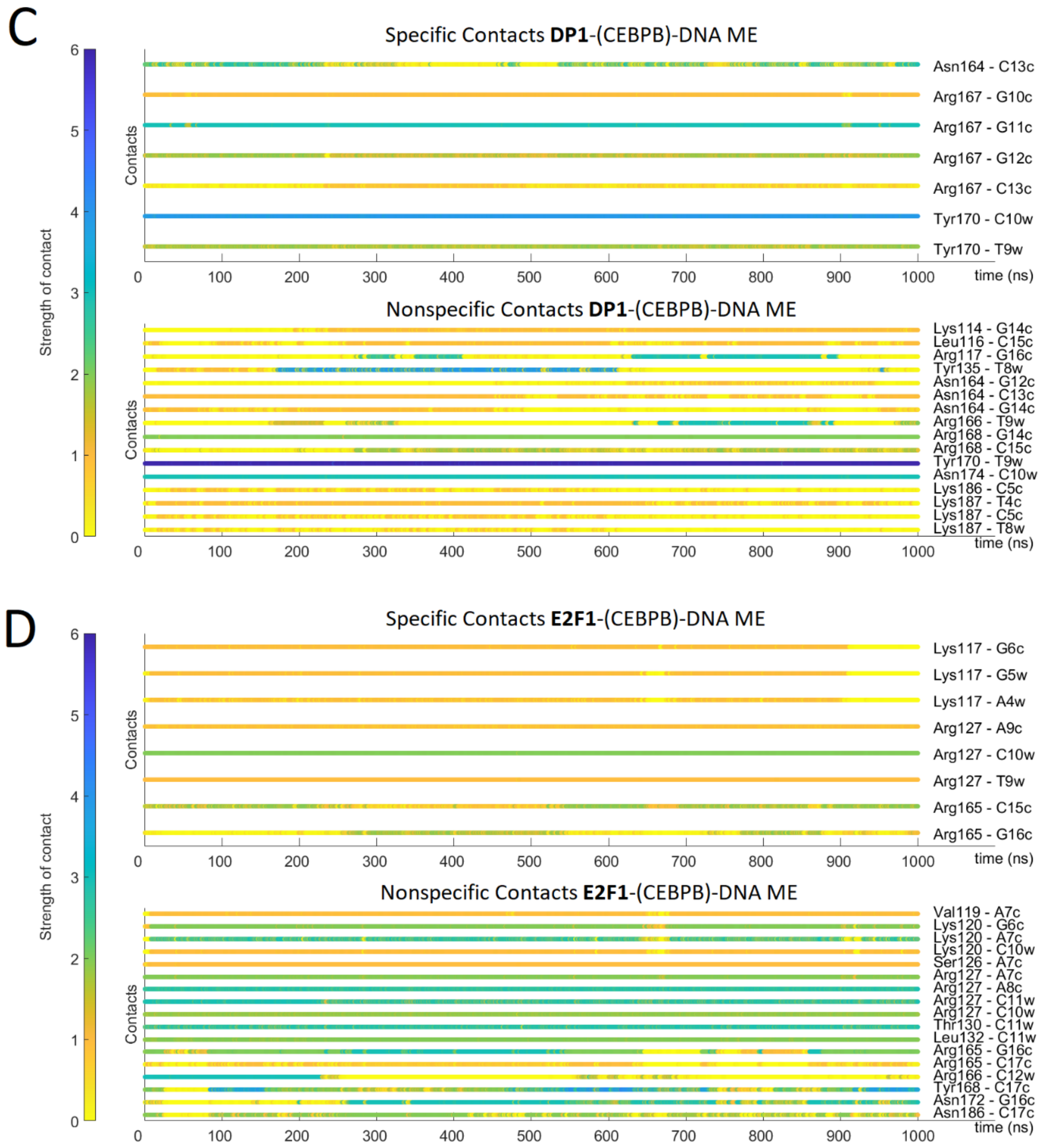


**Figure S8.** Time-evolution of the specific and nonspecific protein-DNA contacts for the E2F1-DP1-CEBPB-DNA system: **A:** WT DP1-DNA; **B:** WT E2F1-DNA; **C:** ME DP1-DNA; **D:** ME E2F1-DNA.

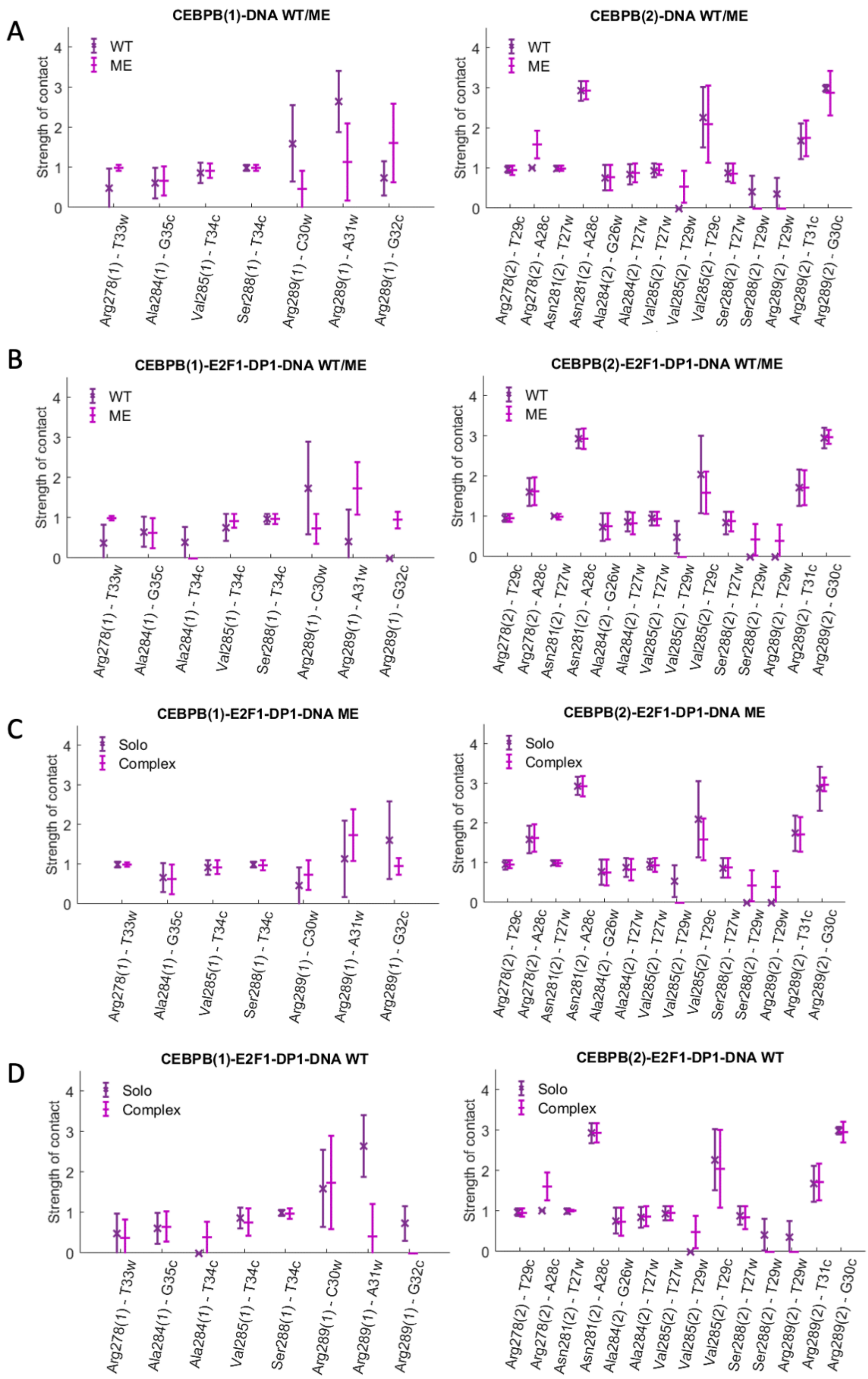


**Figure S9.** Specific contacts between CEBPB-dimer and DNA. The plots show the strength of specific contacts formed by the two CEBPB monomers, marked with (1) and (2), in wild-type (WT) and methylated (ME) cases, when bound alone to DNA (solo) and together with E2F1-DP1 (complex). For the definition of a contact strength see Supplementary Methods.

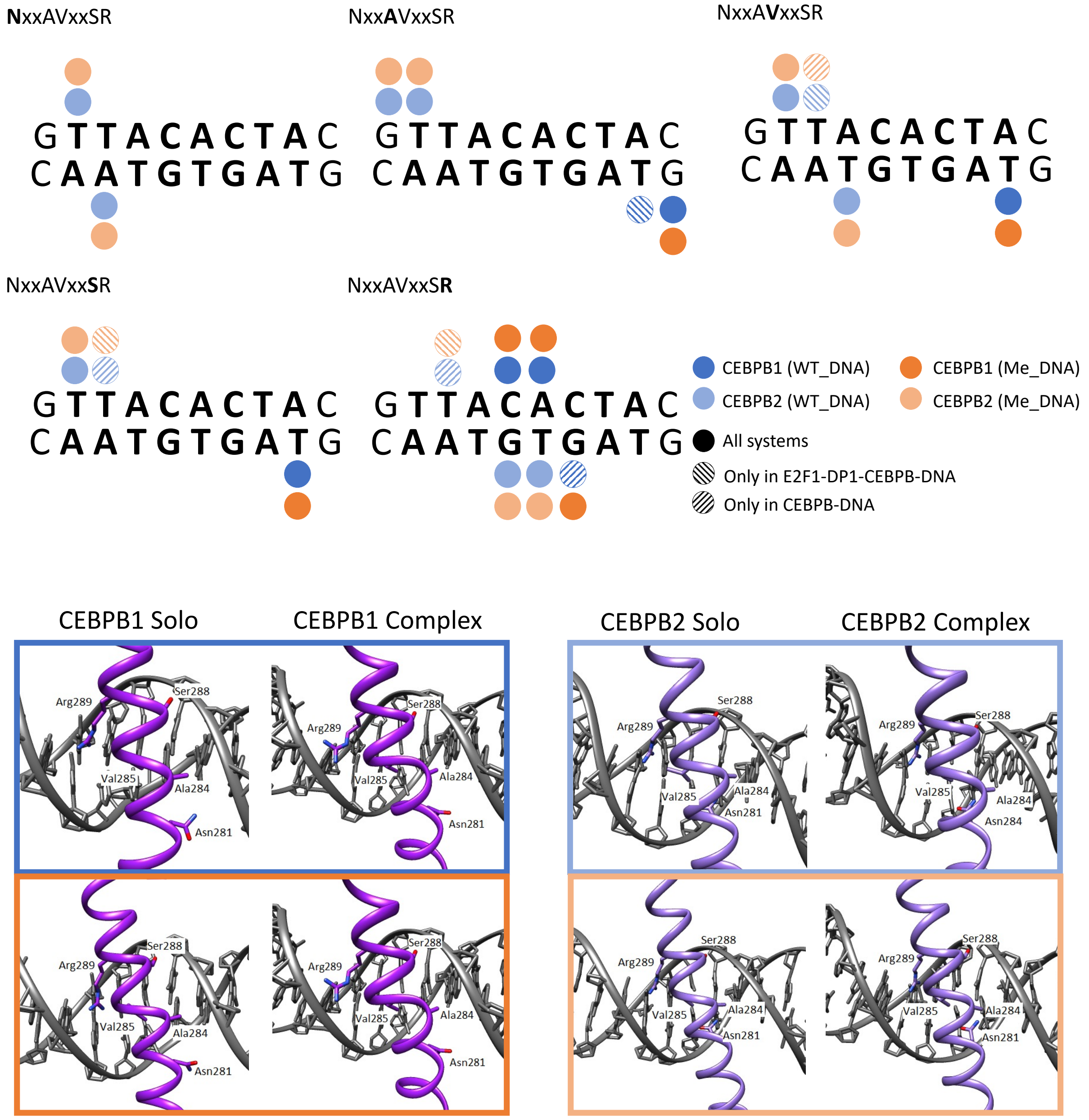


**Figure S10: Above:** Schematic representation of specific protein-DNA contacts exploited by the five-residues-motif (NxxAVxxSR) of CEBPB-dimer in different systems. **Below:** Cartoon representation of the binding orientations and DNA contacts of the CEBPB dimer for WT (blue) and ME (orange) systems, when bound alone to DNA (solo) and together with E2F1-DP1 (complex).

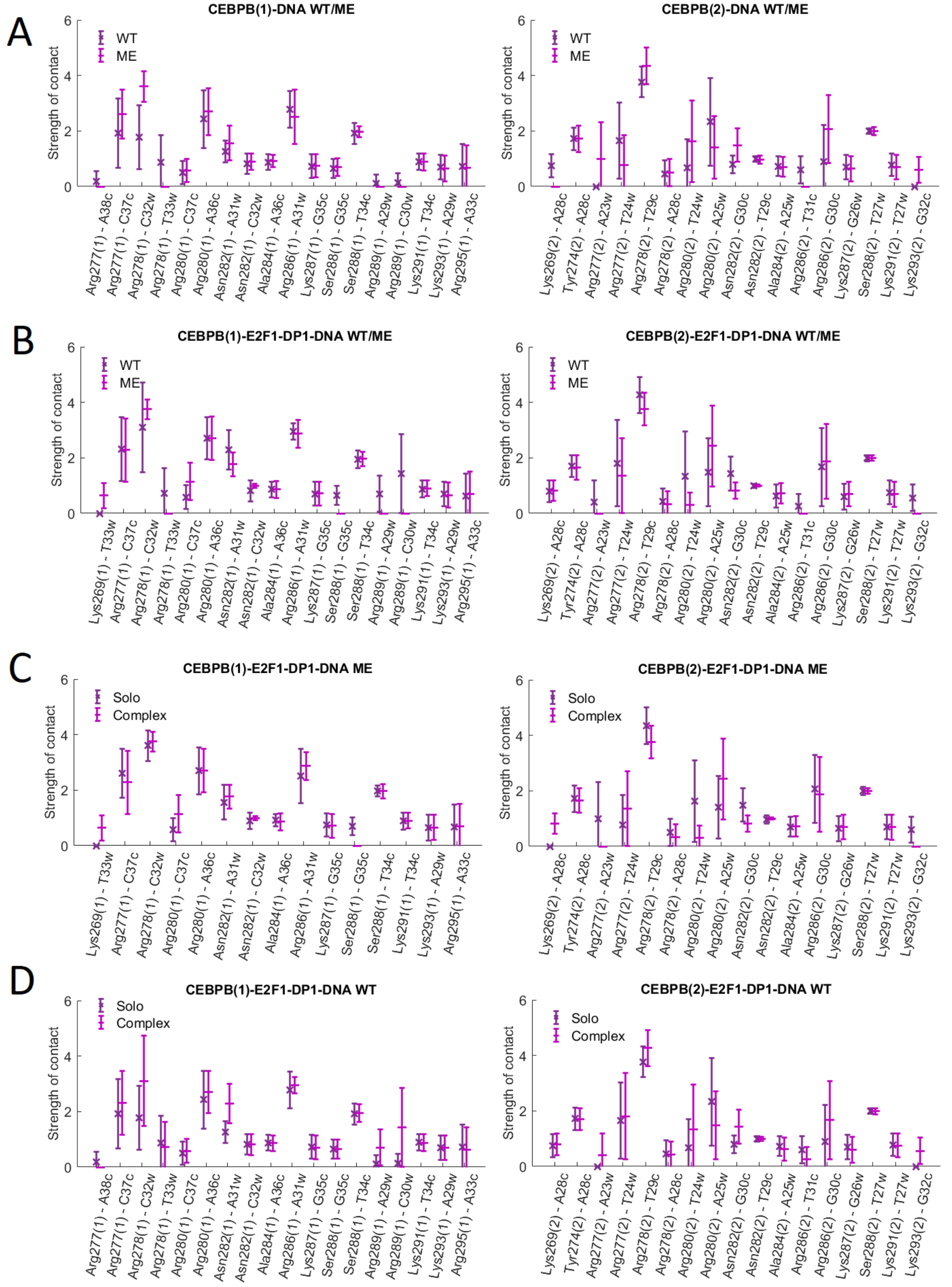


**Figure S11:** Nonspecific contacts between CEBPB-dimer and DNA. The plots show the strength of nonspecific contacts formed by the two CEBPB monomers, marked with (1) and (2), in wild-type (WT) and methylated (ME) cases, when bound alone to DNA (solo) and together with E2F1-DP1 (complex). For the definition of a contact strength see Supplementary Methods.

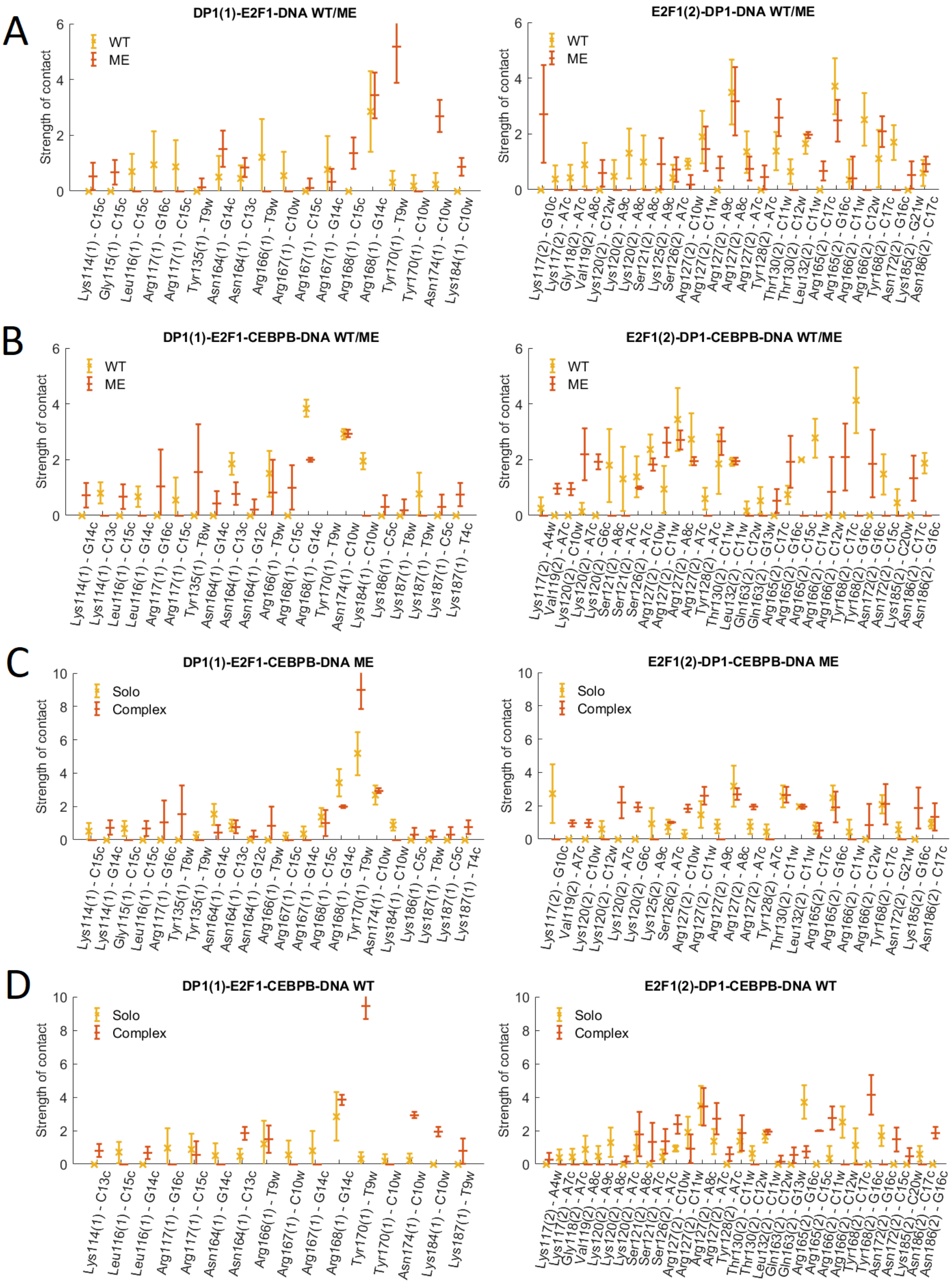


**Figure S12:** Nonspecific contacts between E2F1-DP1-dimer and DNA. The plots show the strength of nonspecific contacts formed by the E2F1 and DP1 monomers, marked with (1) and (2), in wild-type (WT) and methylated (ME) cases, when bound alone to DNA (solo) and together with E2F1-DP1 (complex). For the definition of a contact strength see Supplementary Methods.

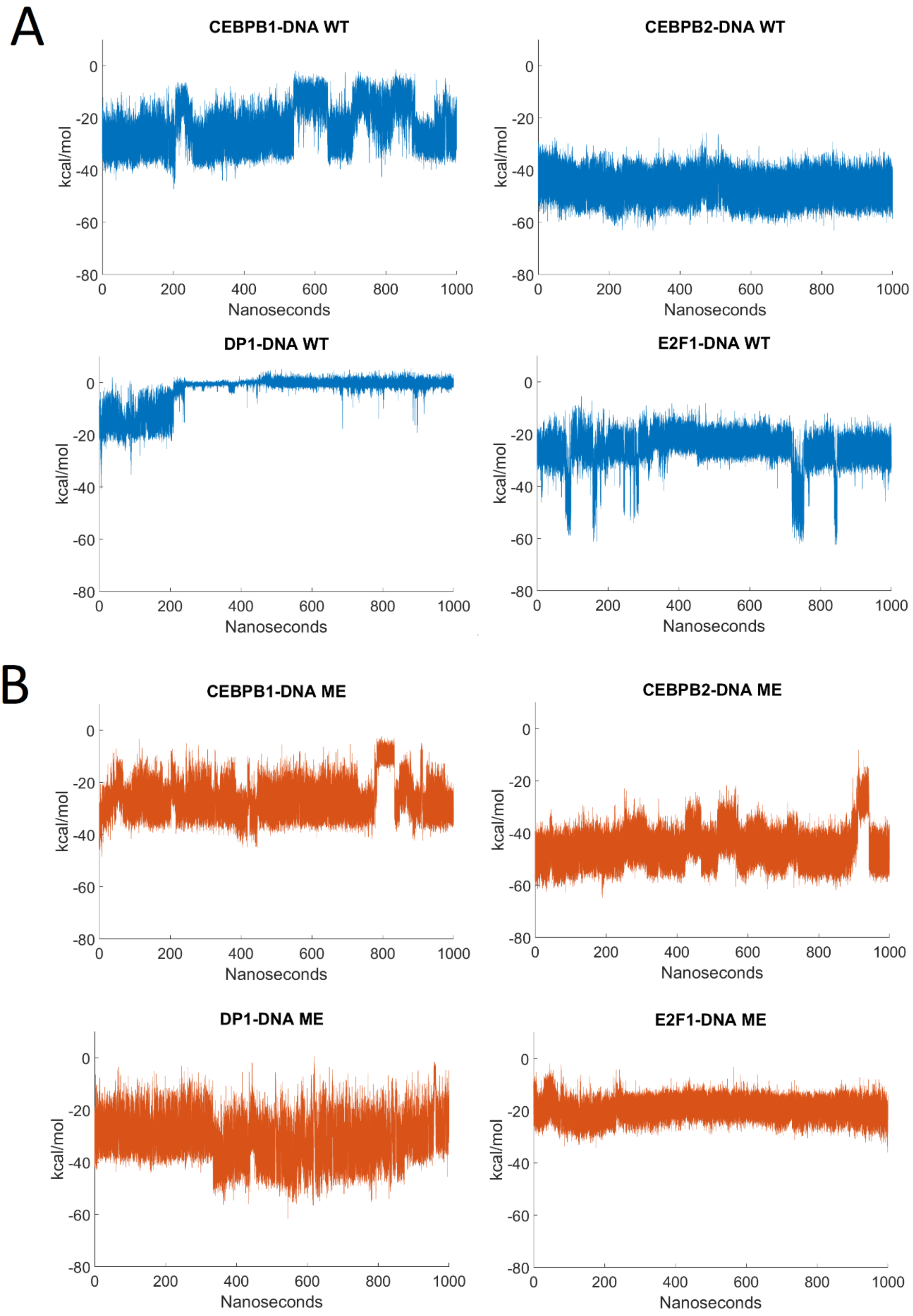


**Figure S13:** Time-evolution of specific interaction energies for the different solo bound TF-DNA complexes for **A:** WT systems (blue) and **B:** ME systems (red).

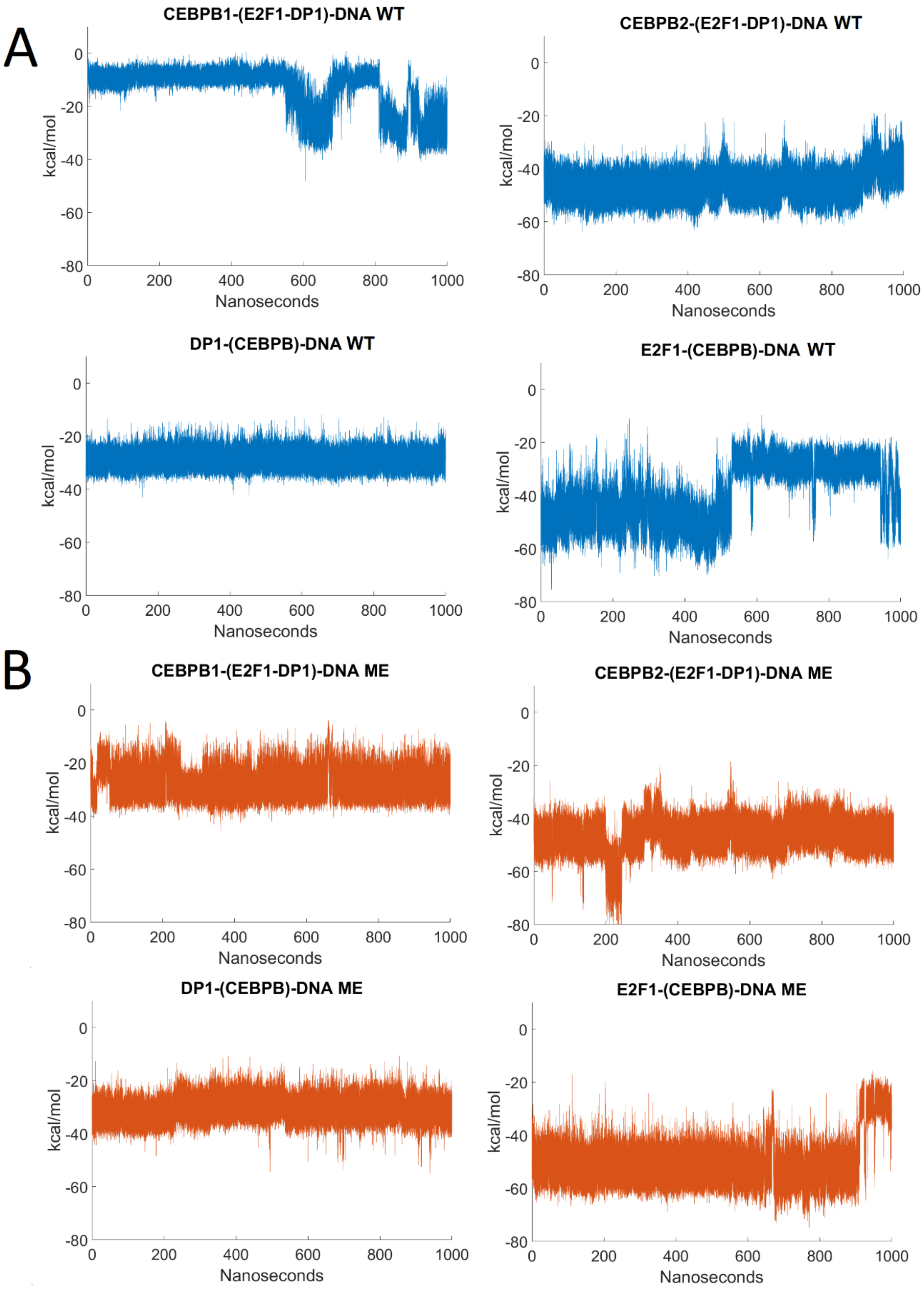


**Figure S14:** Time-evolution of specific interaction energies for the different TF-DNA complexes indicated in parentheses for complete enhanceosome complexes (CEBPB-E2F1-DP1-DNA) for **A:** WT systems (blue) and **B:** ME systems (red).

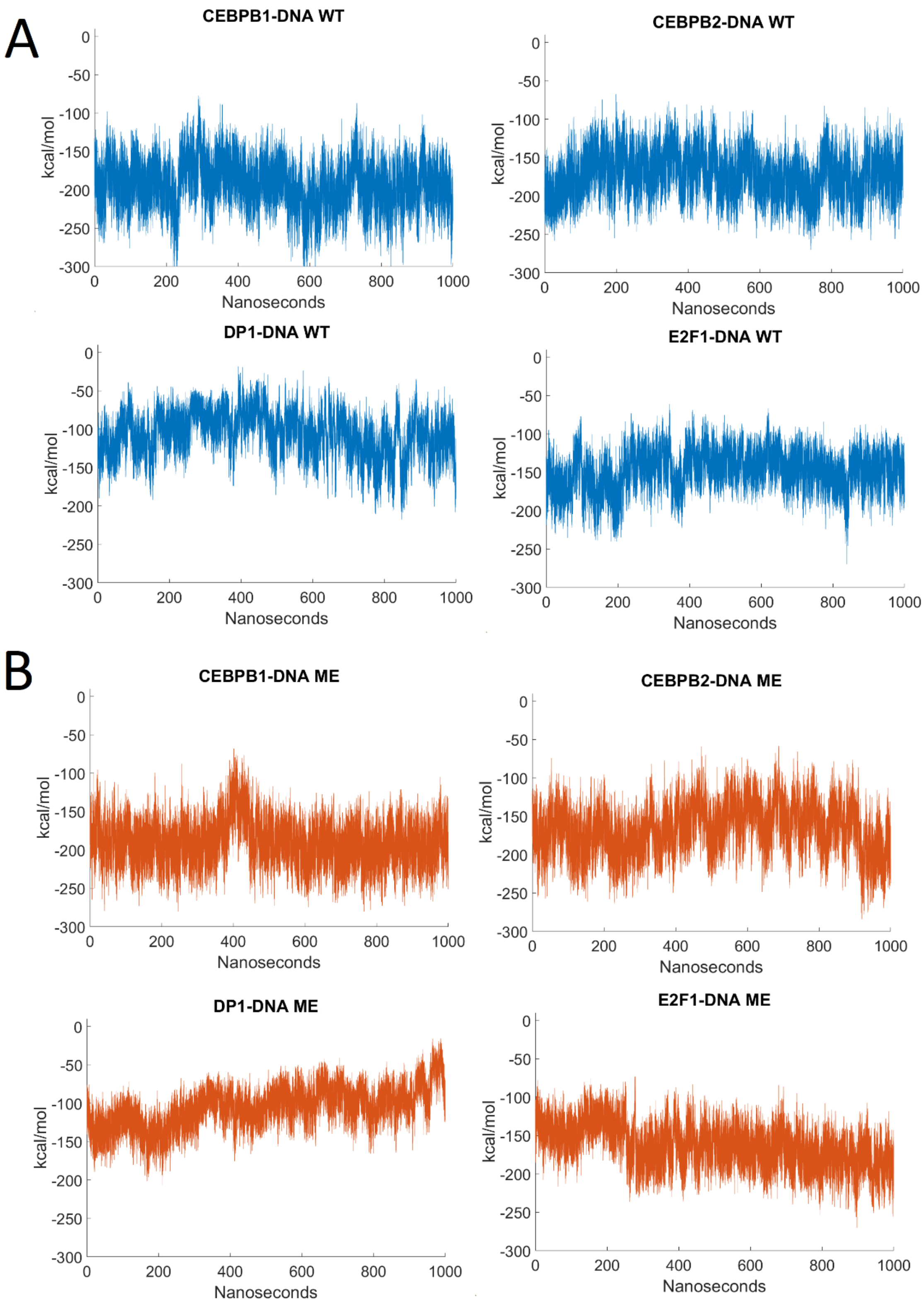


**Figure S15:** Time-evolution of nonspecific interaction energies for the different solo bound TF-DNA complexes for **A:** WT systems (blue) and **B:** ME systems (red).

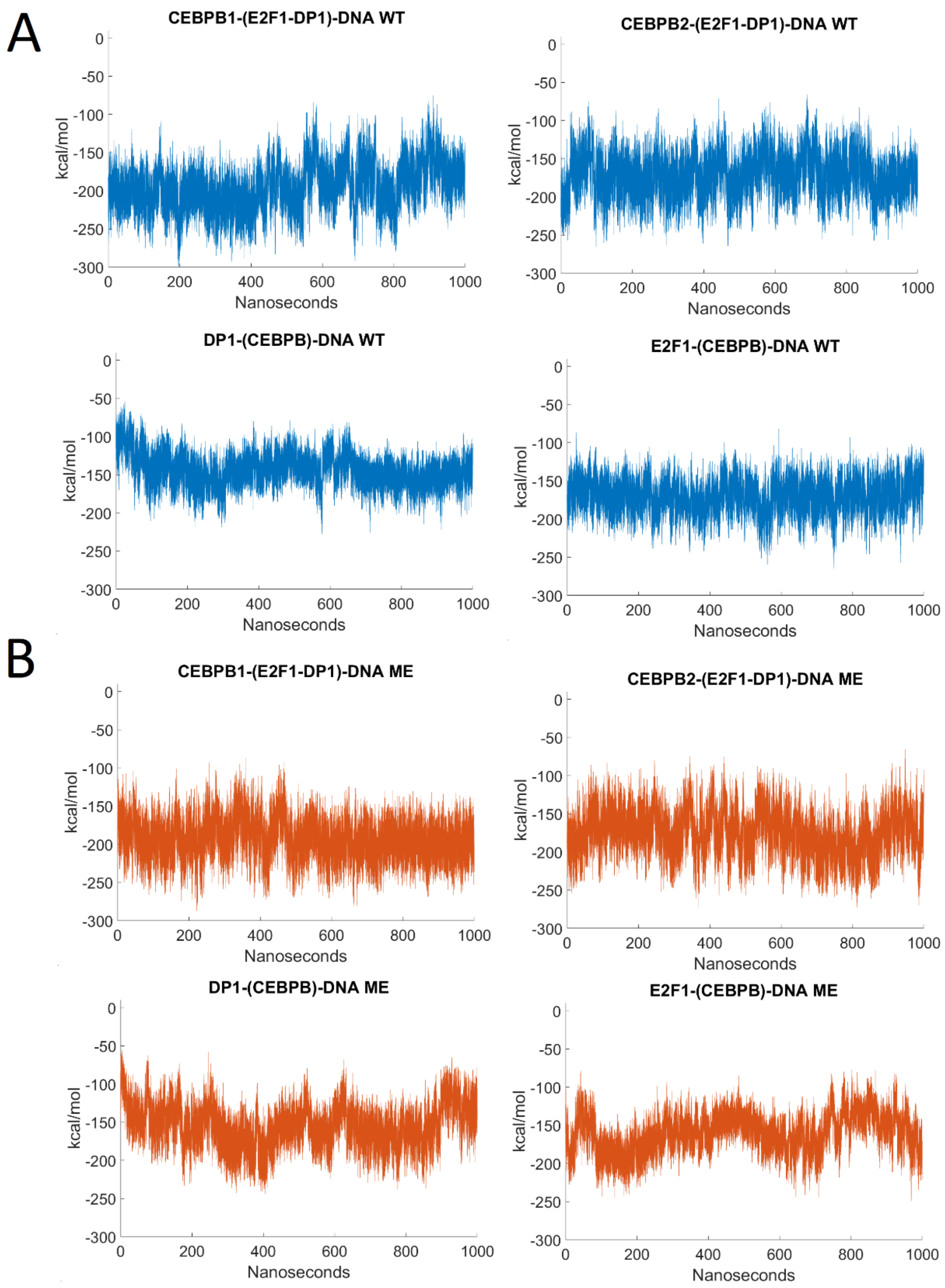


**Figure S16:** Time-evolution of nonspecific interaction energies for the different TF-DNA complexes indicated in parentheses for complete enhanceosome complexes (CEBPB-E2F1-DP1-DNA) for **A:** WT systems (blue) and **B:** ME systems (red).

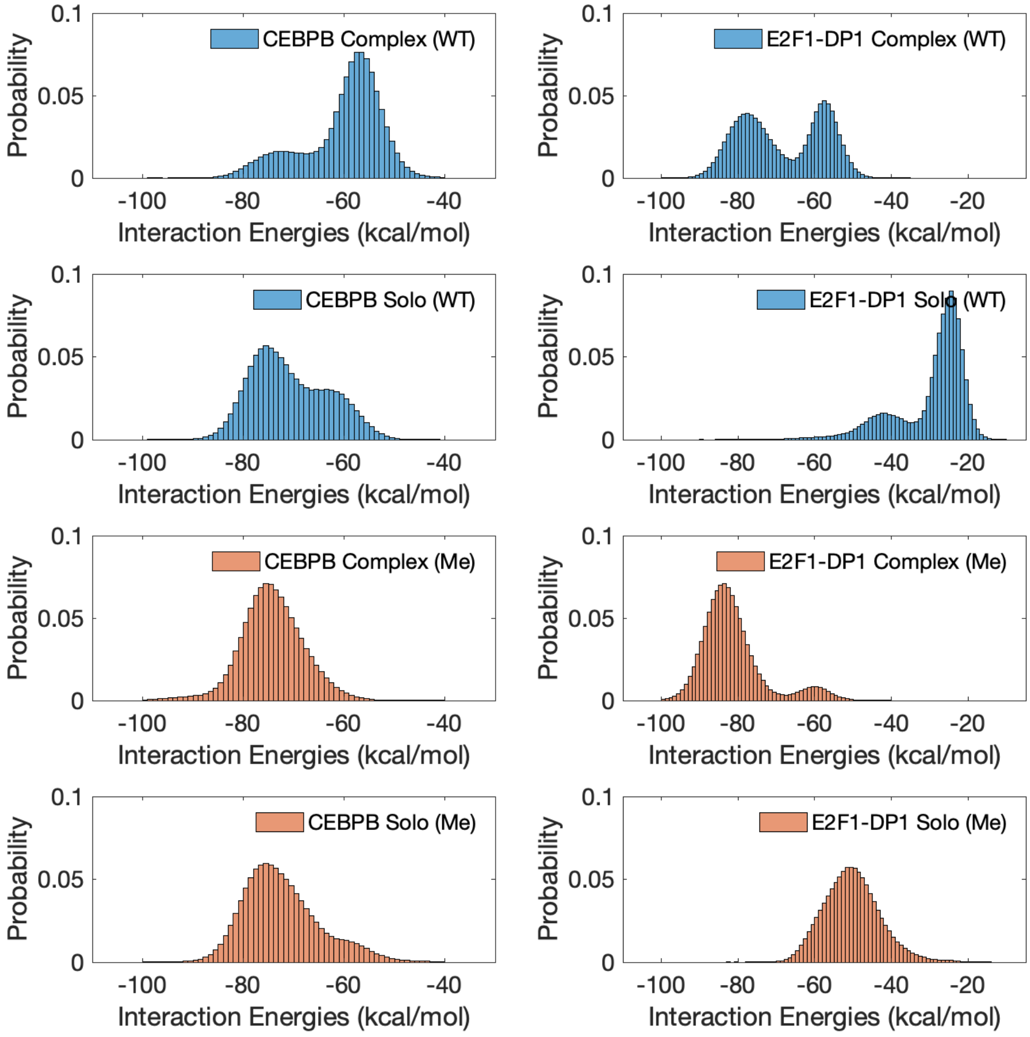


**Figure S17:** Specific contacts interaction energies distributions for the corresponding TF-dimer-DNA complex, were “Solo” corresponds to either CEBPB-DNA or E2F1-DP1-DNA systems, and “Complex” – to CEBPB-E2F1-DP1-DNA systems, the “WT” and “ME” denote wild type and methylated DNA systems.

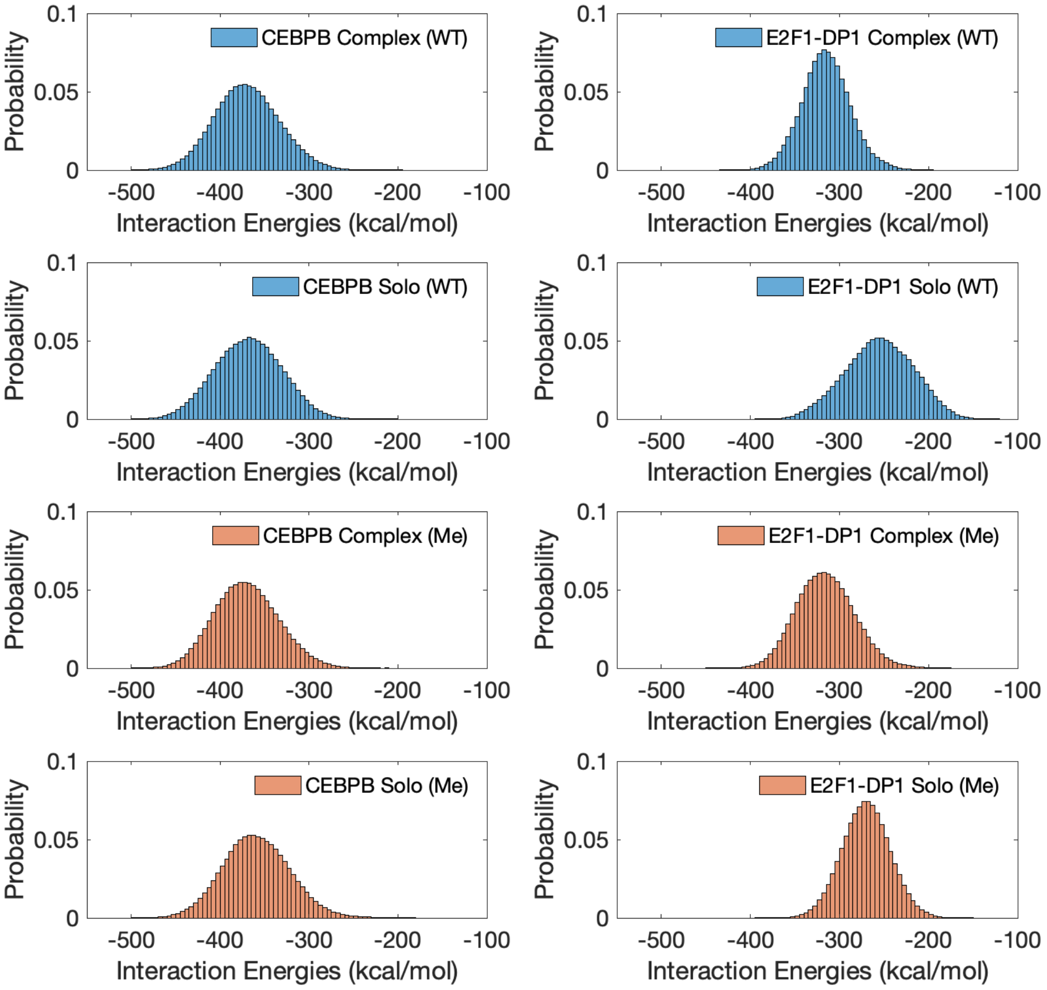


**Figure S18:** Nonspecific contacts interaction energies distributions for the corresponding TF-dimer-DNA complex, were "Solo" corresponds to either CEBPB-DNA or E2F1-DP1-DNA systems, and "Complex" – to CEBPB-E2F1-DP1-DNA systems, the "WT" and "ME" denote wild type and methylated DNA systems.

**Table S1:** Protein-DNA binding energies (kcal/mol) calculated according the MM(GB/PB)SA approach for wild-type (WT) and methylated (ME) systems. For E2F1-DP1-CEBPB-DNA trajectories, the provided interaction energies are for the protein dimer in bold.

| | **CEBPB-DNA** | **E2F1-DP1-DNA** | E2F1-DP1-**CEBPB-DNA** | CEBPB-**E2F1-DP1-DNA** |
|---|---|---|---|---|
| | | | **MMGBSA** | |
| **WT** | -276.5±14.8 | -181.1±15.1 | -276.0±16.1 | -215.4±12.6 |
| **ME** | -277.6±15.9 | -194.6±15.5 | -281.4±13.3 | -214.2±17.0 |
| | | | **MMPBSA** | |
| **WT** | -285.1±15.7 | -204.2±16.1 | -281.5±15.0 | -225.4±14.1 |
| **ME** | -285.5±16.1 | -204.4±16.2 | -289.7±14.4 | -220.5±17.8 |

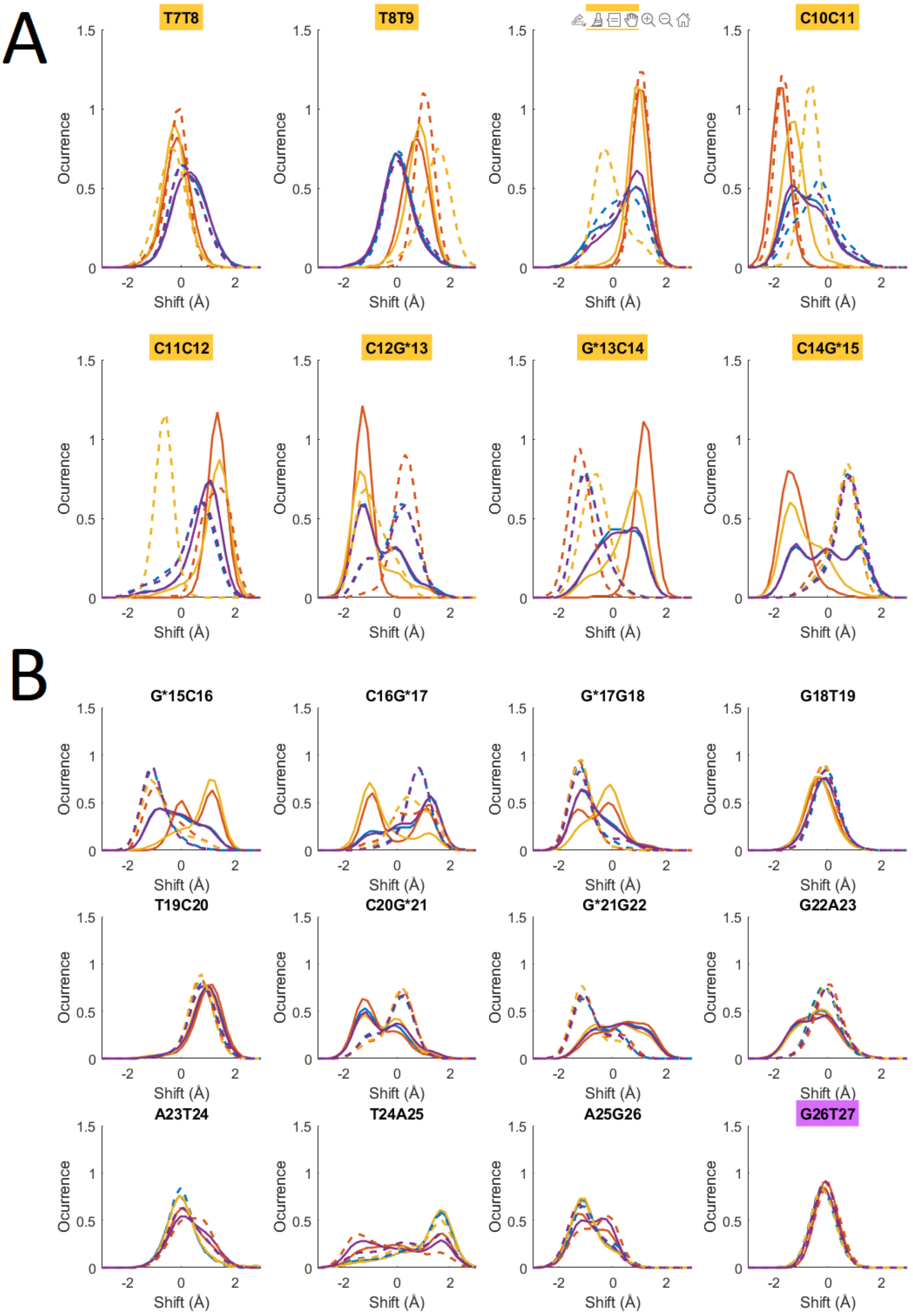

A
T7T8
T8T9
C10C11
C11C12
C12G*13
G*13C14
C14G*15
B
G*15C16
C16G*17
G*17G18
G18T19
T19C20
C20G*21
G*21G22
G22A23
A23T24
T24A25
A25G26
G26T27
Ocurrence
Shift (Å)

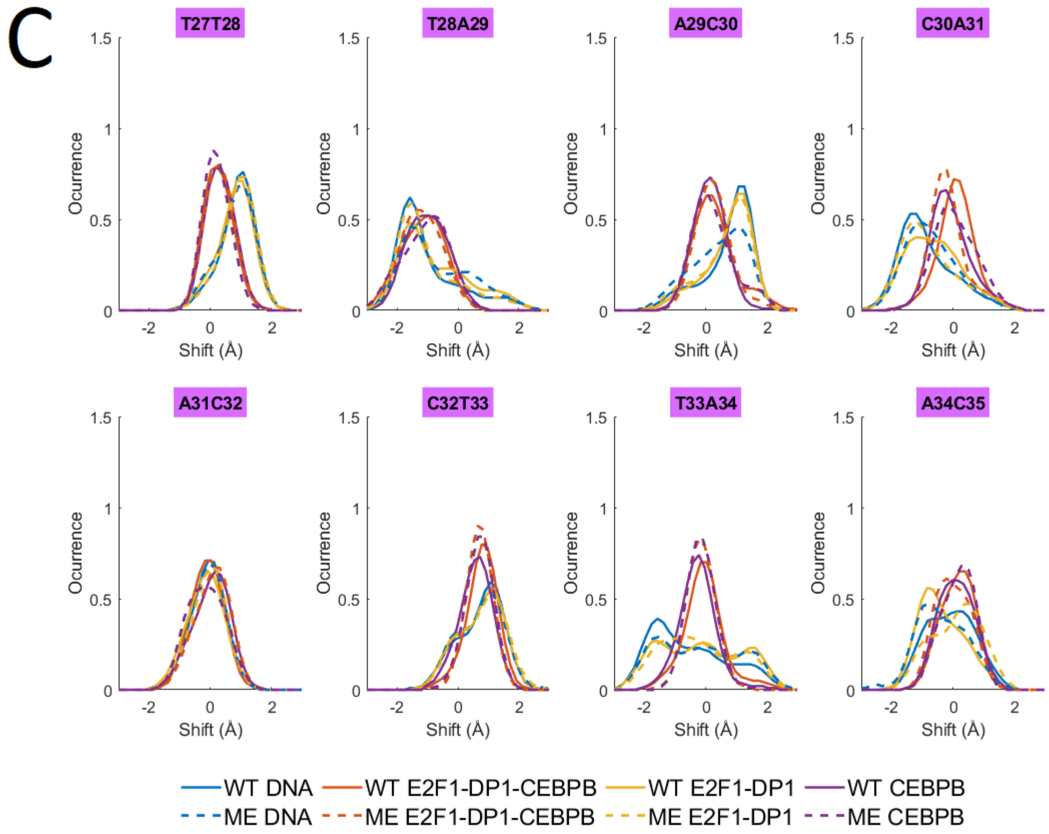


**Figure S19:** Normalized b.p. step shift distributions for DNA alone and in complex with either one (CEBPB or E2F1-DP1 TFs dimer) or both for **A:** the E2F1-DP1 binding site; **B:** the linker region; **C:** the CEBPB binding site.

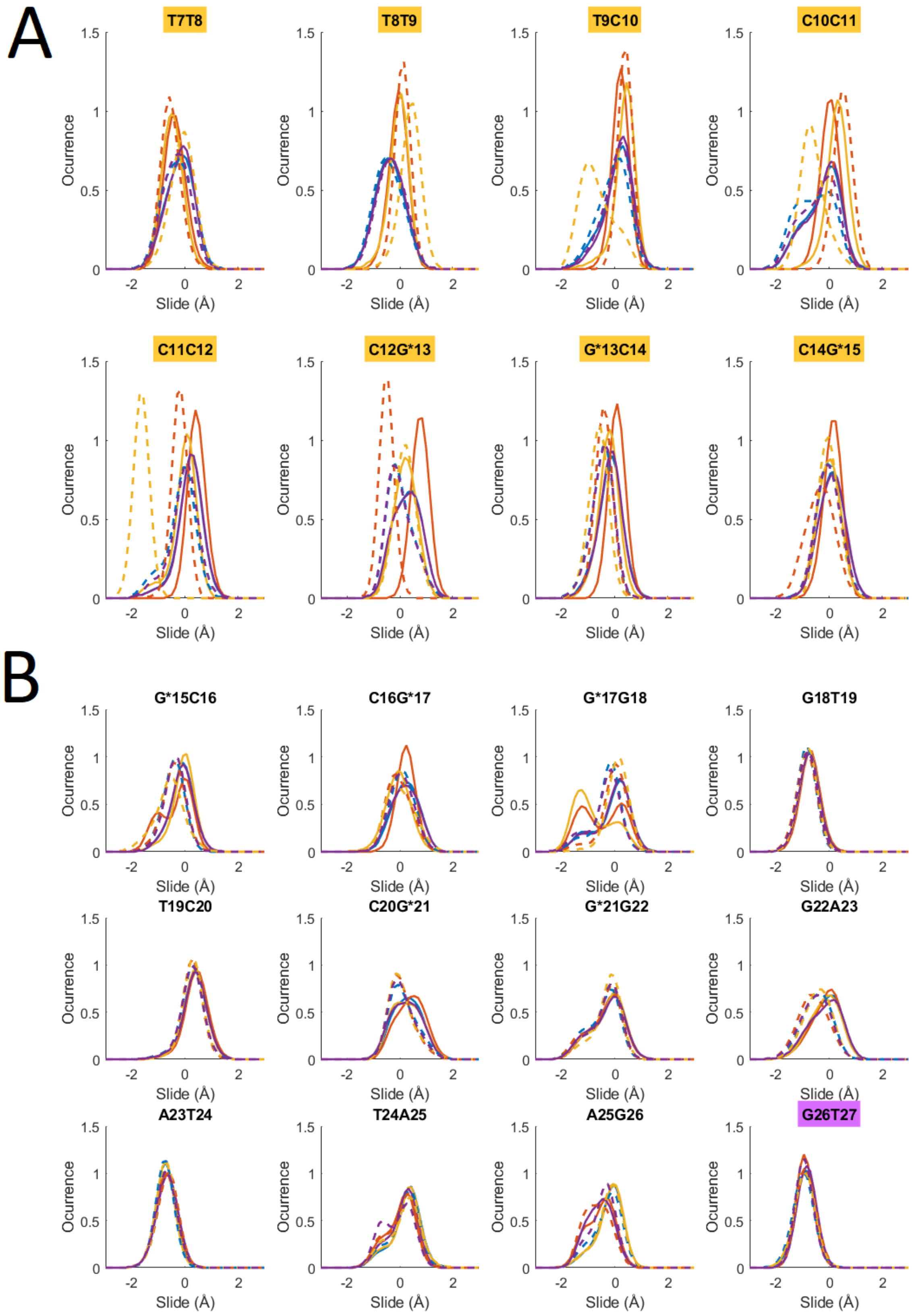
A
T7T8
T8T9
T9C10
C10C11
C11C12
C12G*13
G*13C14
C14G*15
B
G*15C16
C16G*17
G*17G18
G18T19
T19C20
C20G*21
G*21G22
G22A23
A23T24
T24A25
A25G26
G26T27
Ocurrence
Slide (Å)
1.5
1
0.5
0
-2
0
2

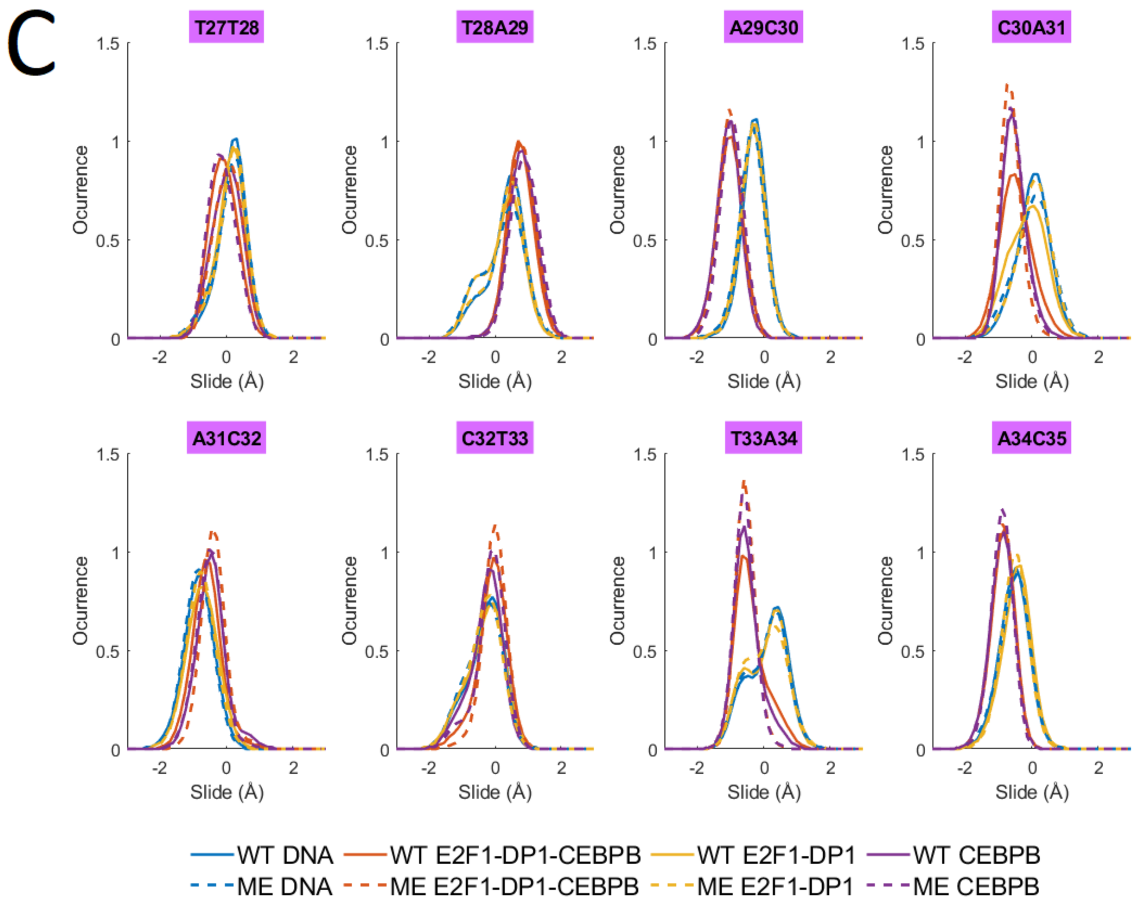


**Figure S20:** Normalized b.p. step slide distributions for DNA alone and in complex with either one (CEBPB or E2F1-DP1 TFs dimer) or both for **A:** the E2F1-DP1 binding site; **B:** the linker region; **C:** the CEBPB binding site.

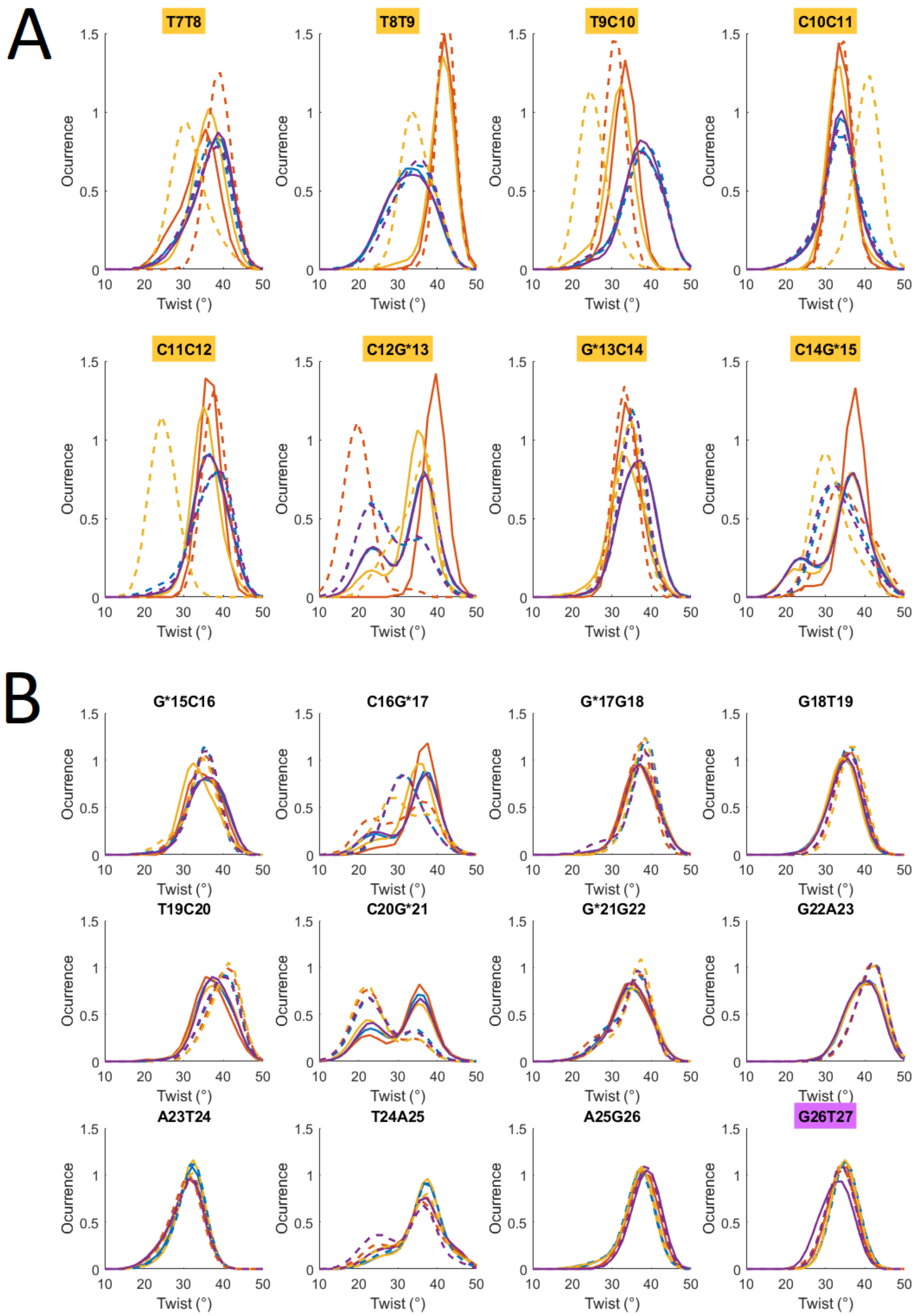

A
T7T8
T8T9
T9C10
C10C11
C11C12
C12G*13
G*13C14
C14G*15
B
G*15C16
C16G*17
G*17G18
G18T19
T19C20
C20G*21
G*21G22
G22A23
A23T24
T24A25
A25G26
G26T27
Ocurrence
Twist (°)

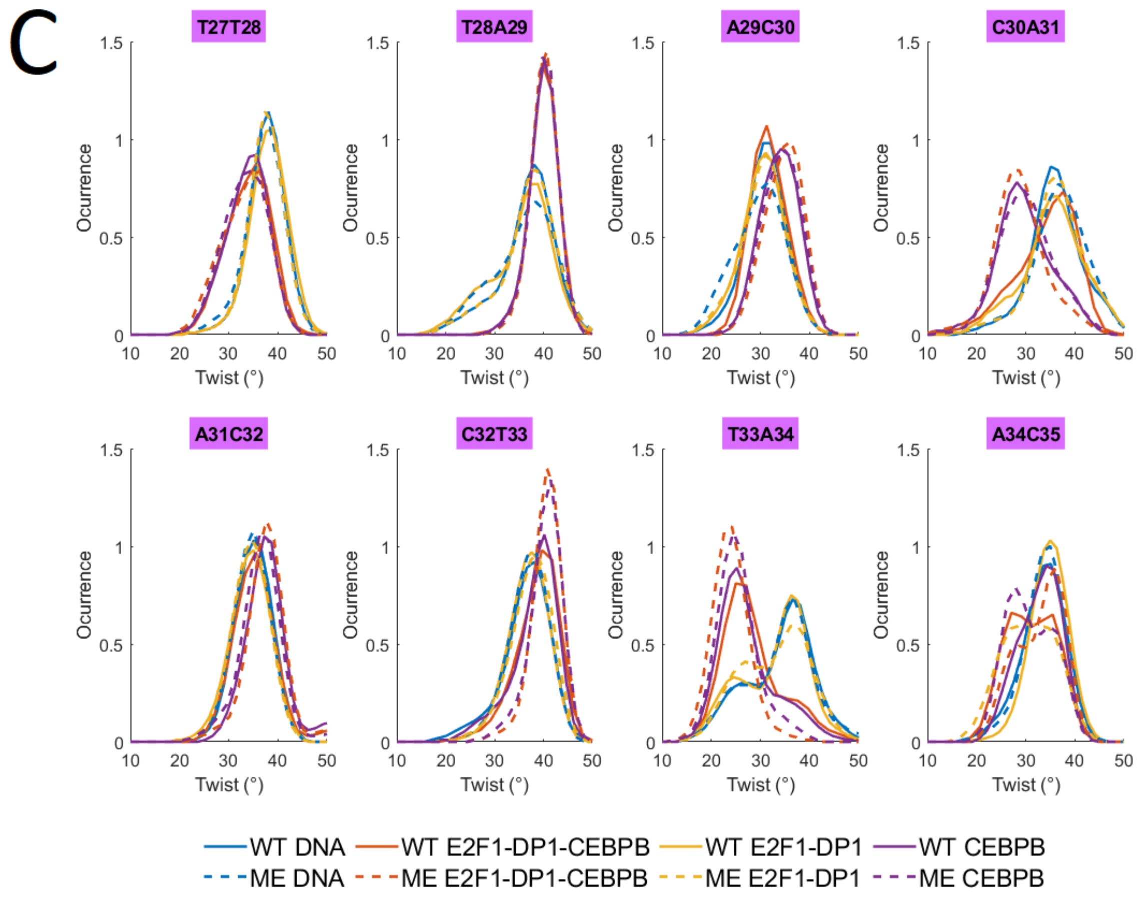


**Figure S21:** Normalized b.p. step twist distributions for DNA alone and in complex with either one (CEBPB or E2F1-DP1 TFs dimer) or both for **A:** the E2F1-DP1 binding site; **B:** the linker region; **C:** the CEBPB binding site.

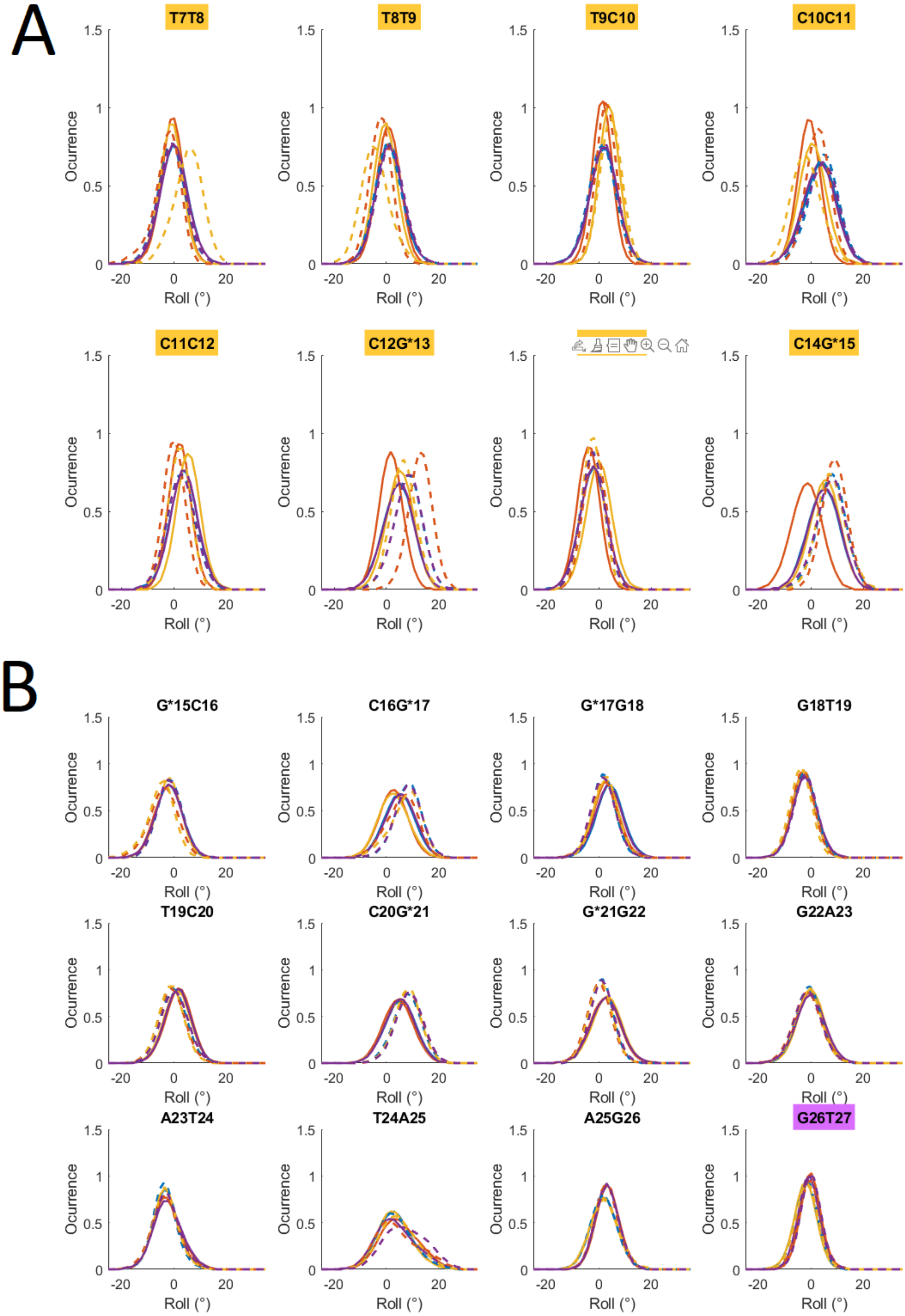
A
T7T8
T8T9
T9C10
C10C11
C11C12
C12G*13
C14G*15
B
G*15C16
C16G*17
G*17G18
G18T19
T19C20
C20G*21
G*21G22
G22A23
A23T24
T24A25
A25G26
G26T27
Ocurrence
Roll (°)

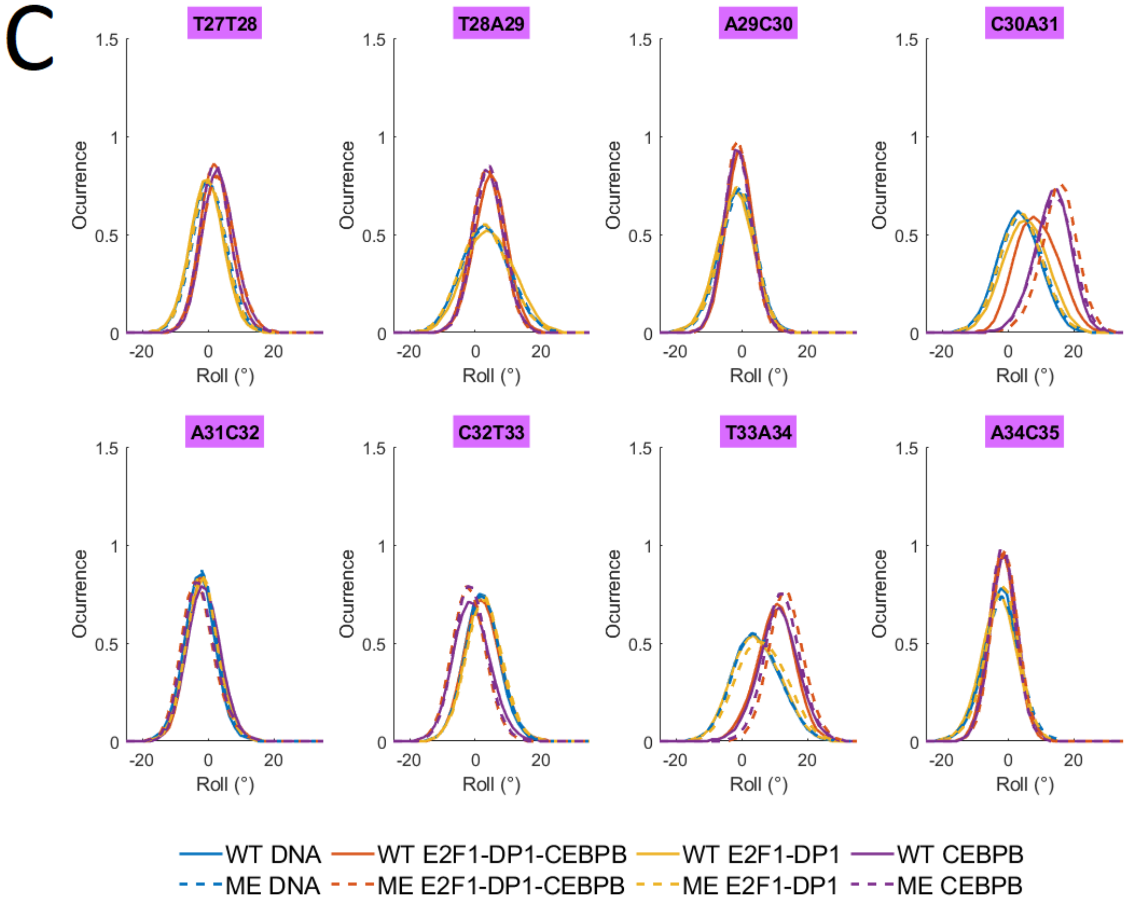


**Figure S22:** Normalized b.p. step roll distributions for DNA alone and in complex with either one (CEBPB or E2F1-DP1 TFs dimer) or both for **A:** the E2F1-DP1 binding site; **B:** the linker region; **C:** the CEBPB binding site.

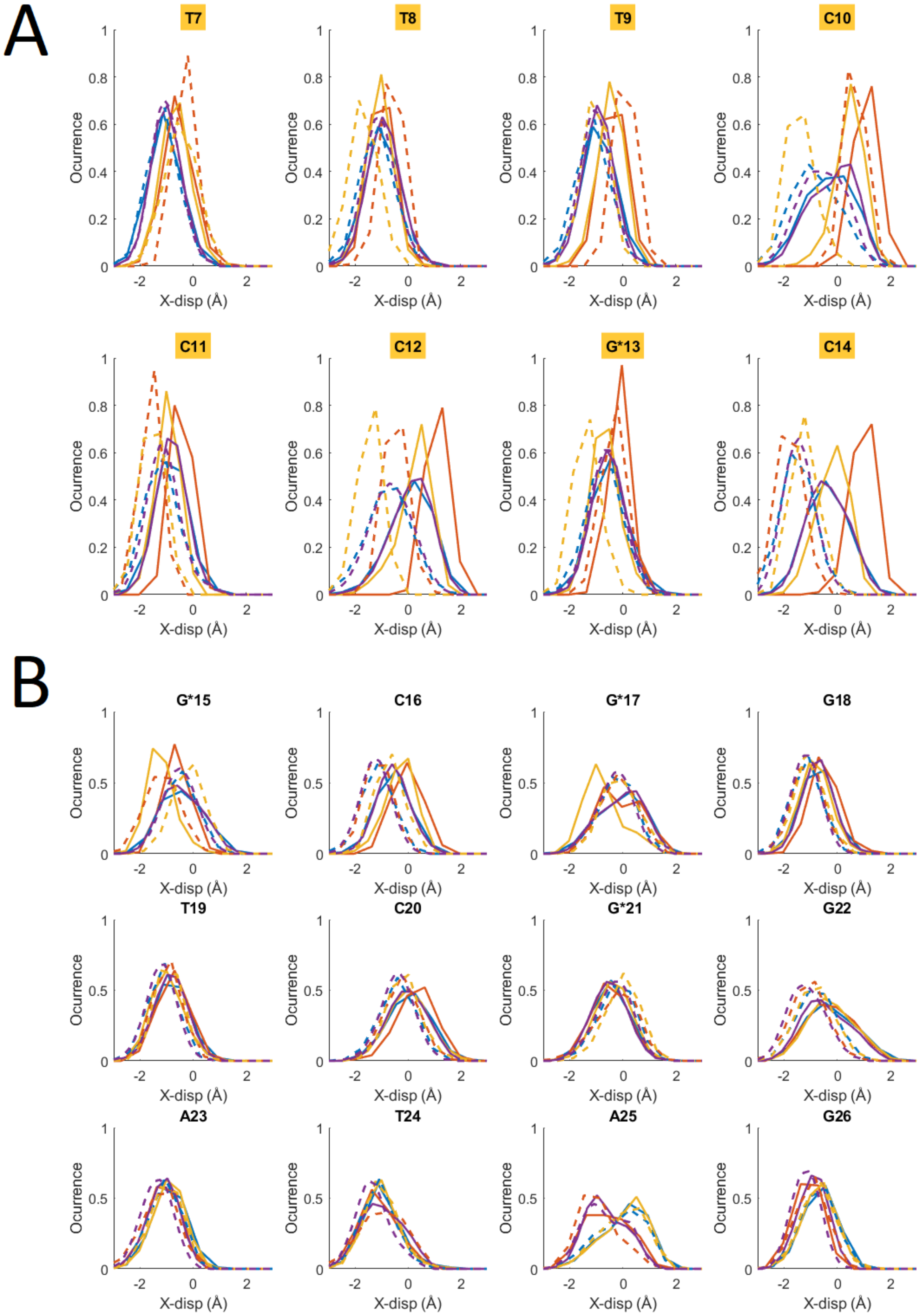
A
T7
T8
T9
C10
C11
C12
G*13
C14
B
G*15
C16
G*17
G18
T19
C20
G*21
G22
A23
T24
A25
G26
Ocurrence
X-disp (Å)

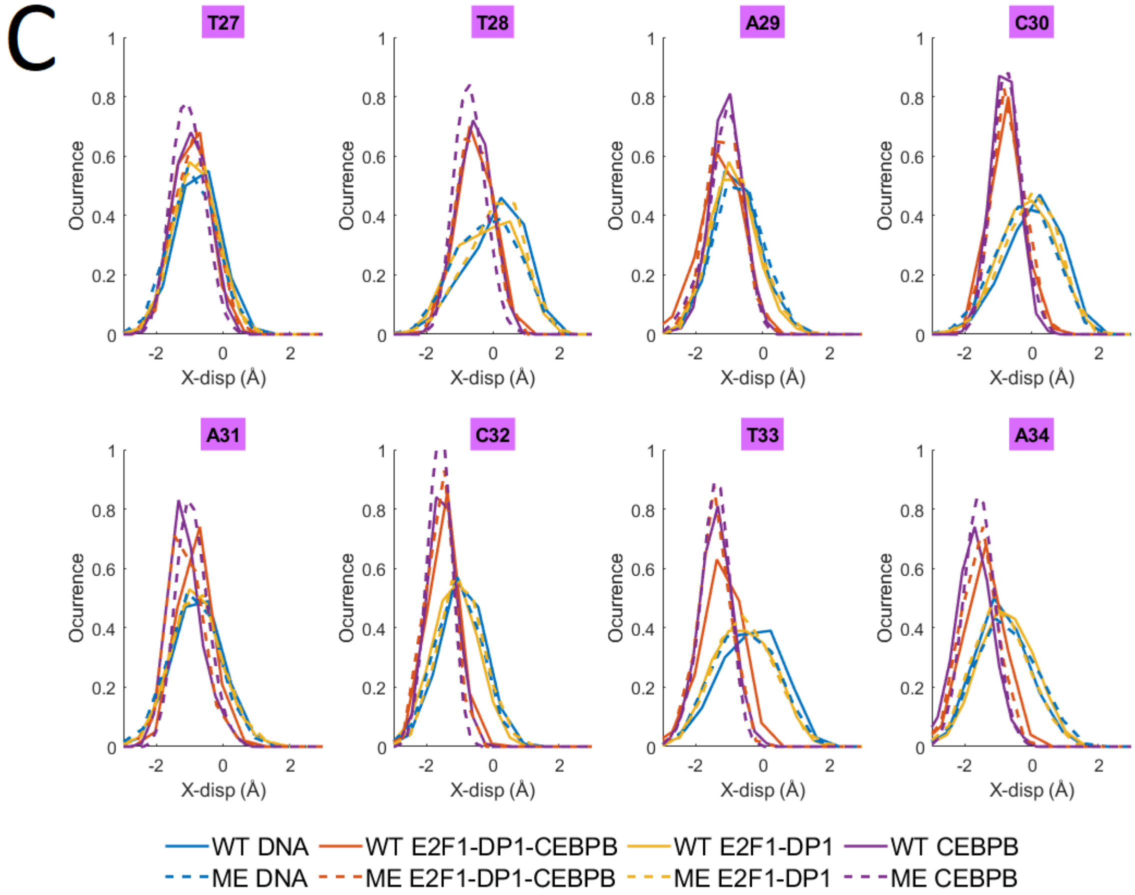


**Figure S23:** Normalized b.p. x-displacement distributions for DNA alone and in complex with either one (CEBPB or E2F1-DP1 TFs dimer) or both for **A:** the E2F1-DP1 binding site; **B:** the linker region; **C:** the CEBPB binding site.

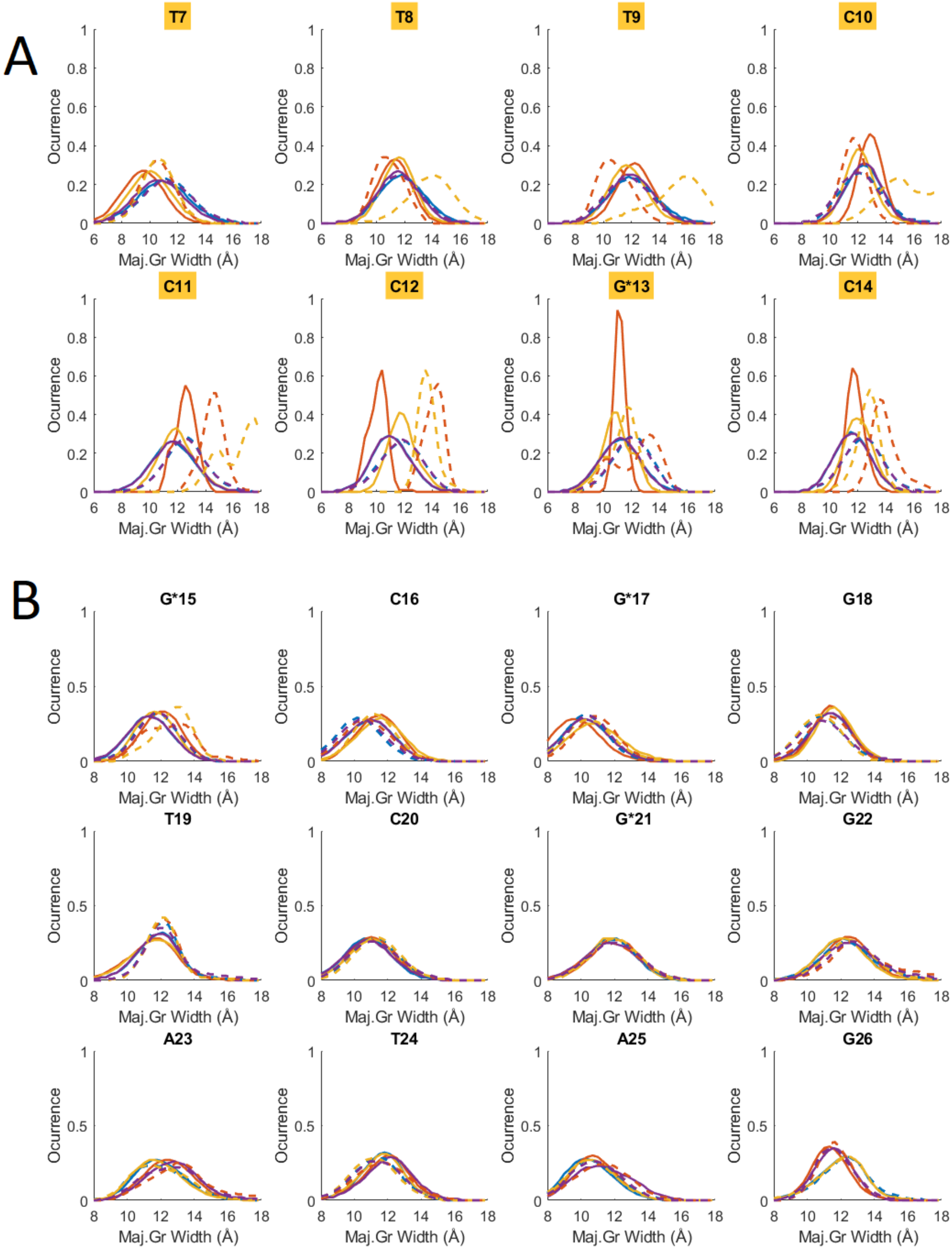
A
T7
T8
T9
C10
C11
C12
G*13
C14
B
G*15
C16
G*17
G18
T19
C20
G*21
G22
A23
T24
A25
G26
Ocurrence
Maj.Gr Width (Å)

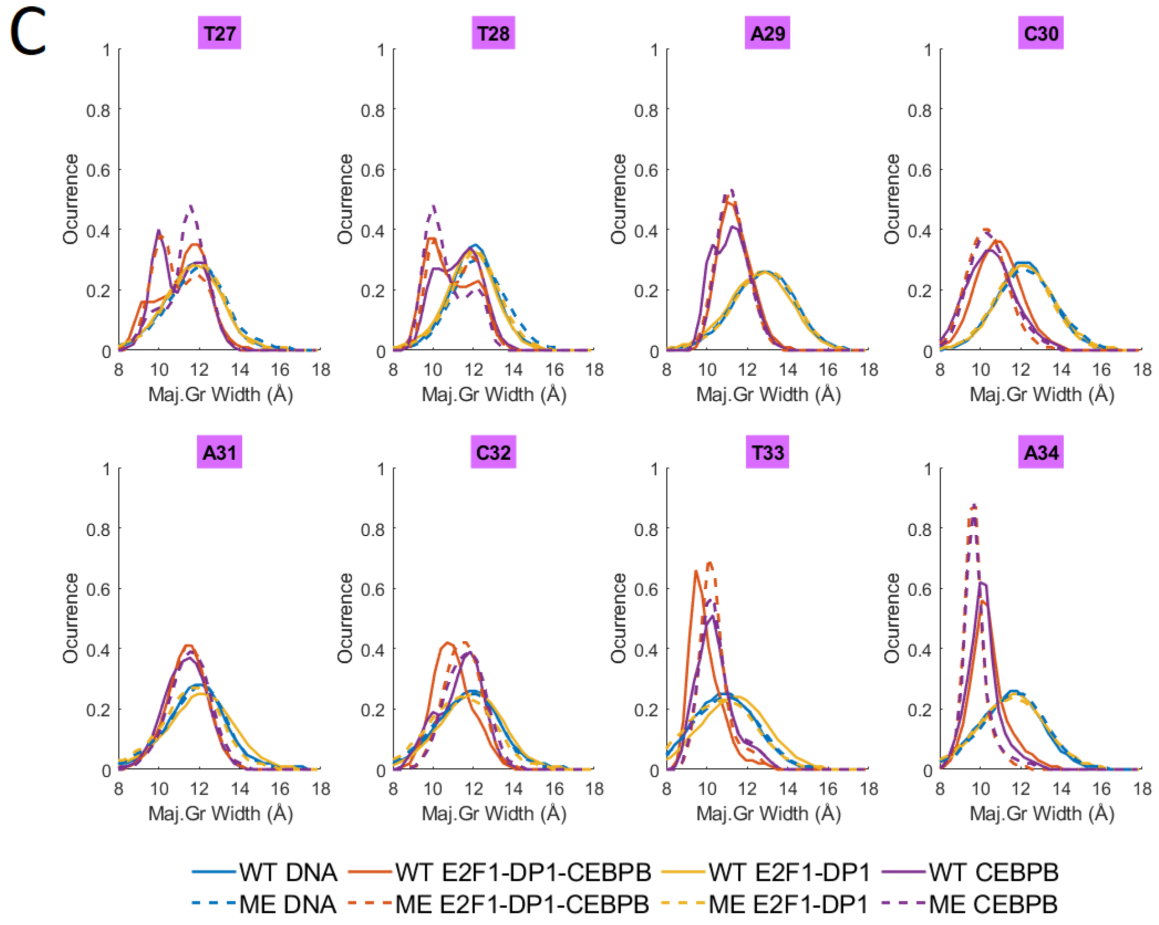


**Figure S24:** Normalized major groove width distributions for DNA alone and in complex with either one (CEBPB or E2F1-DP1 TFs dimer) or both for **A:** the E2F1-DP1 binding site; **B:** the linker region; **C:** the CEBPB binding site.

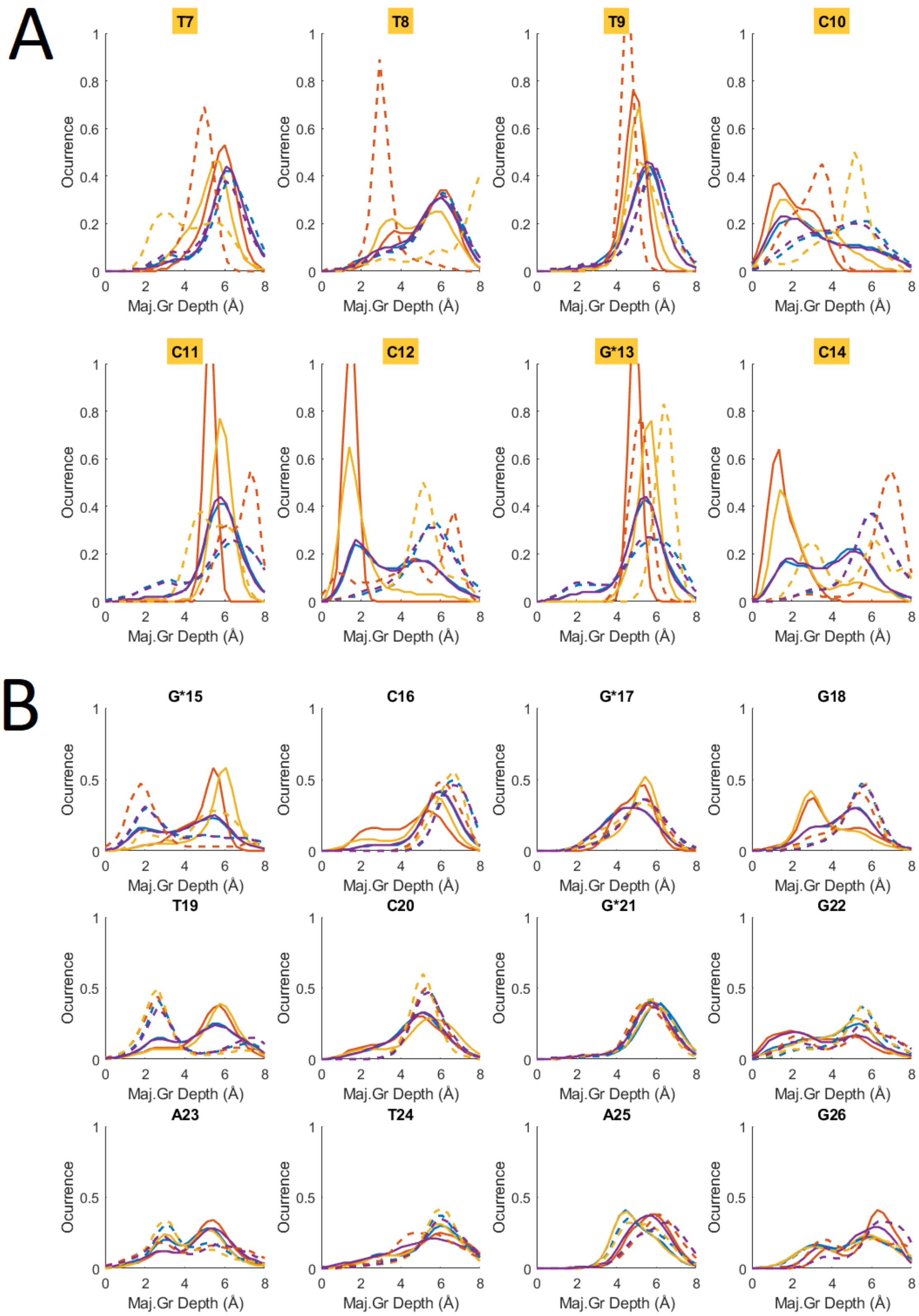

A
T7
T8
T9
C10
C11
C12
G*13
C14
B
G*15
C16
G*17
G18
T19
C20
G*21
G22
A23
T24
A25
G26
Ocurrence
Maj.Gr Depth (Å)

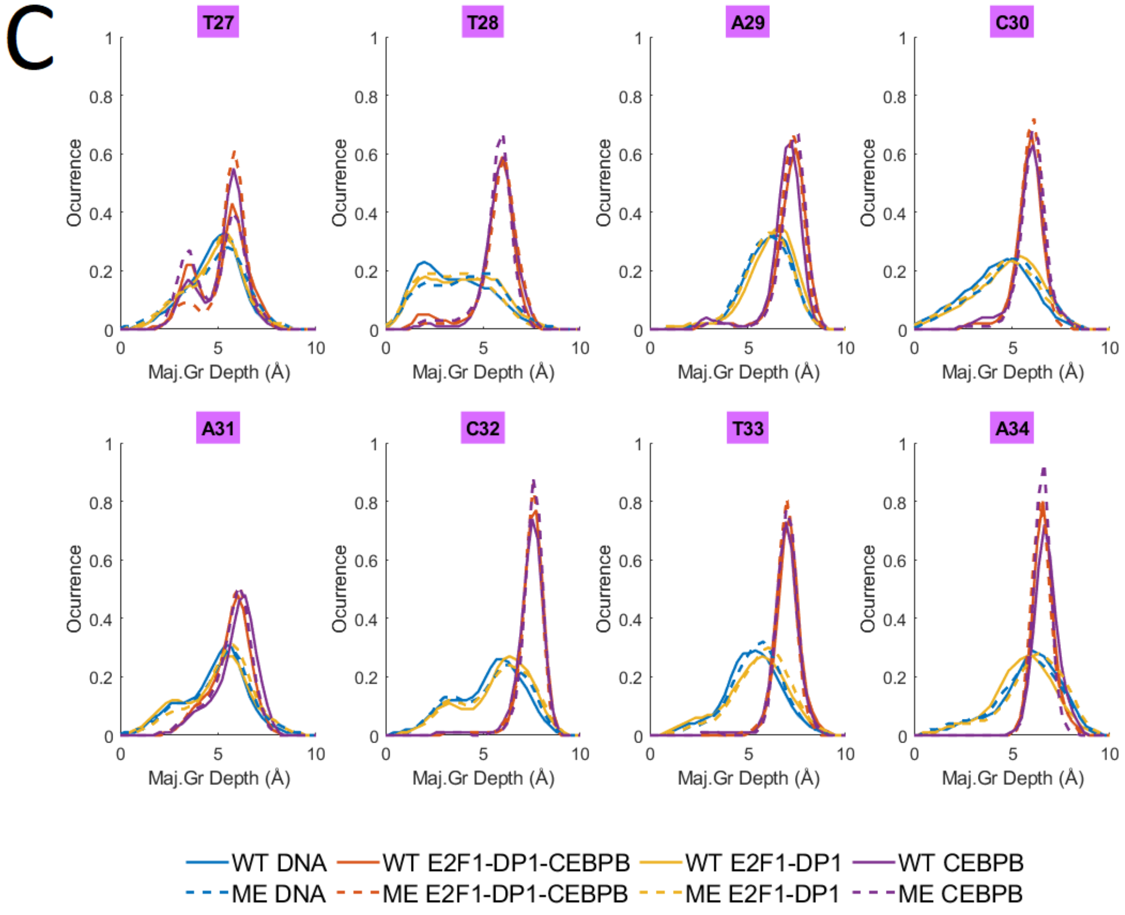


**Figure S25:** Normalized major groove depth distributions for DNA alone and in complex with either one (CEBPB or E2F1-DP1 TFs dimer) or both for **A:** the E2F1-DP1 binding site; **B:** the linker region; **C:** the CEBPB binding site.

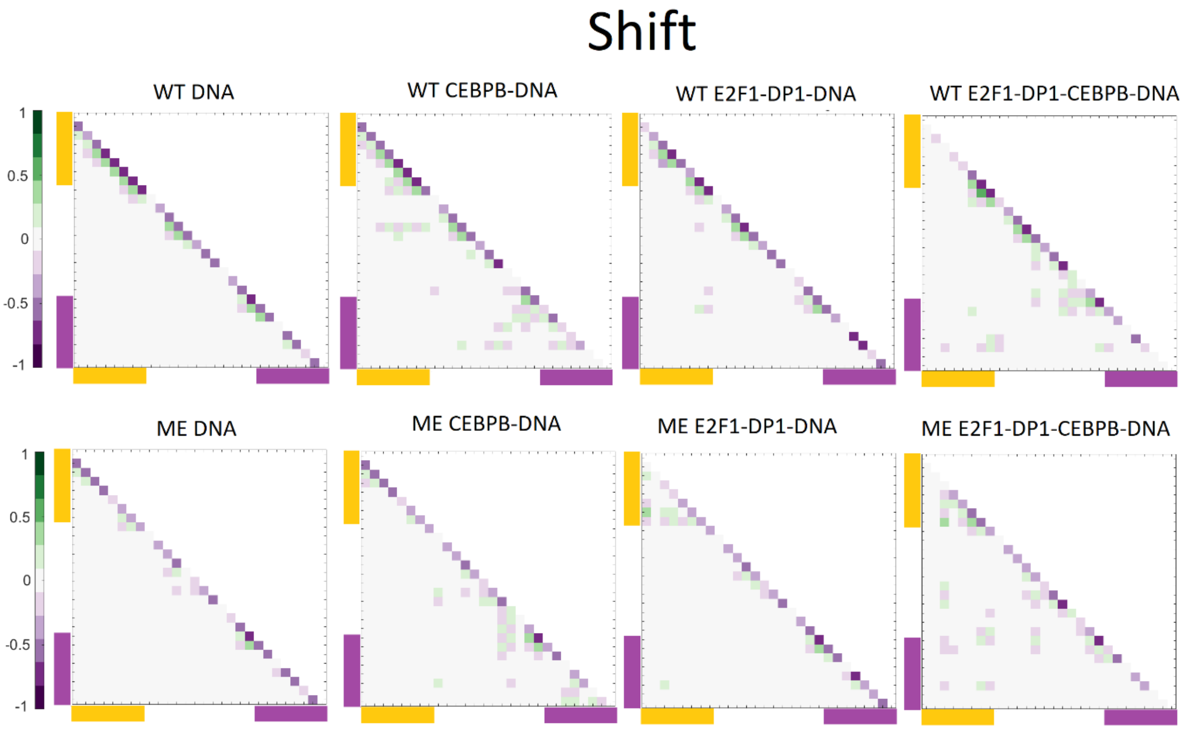


**Figure S26:** Time-delayed correlation coefficient maps for b.p. step shift, with a maximum allowed time lag of 10 ns along the WT and ME sequences for DNA alone and in complex with either CEBPB/E2F1-DP1 dimers, or with both TFs dimers CEBPB-E2F1-DP1. Correlation coefficients are calculated for the entire (0.1–1.1 μs) trajectories. In all panels, the E2F1-DP1 response element is marked with yellow, the CEBPB response element – with magenta. The maximum correlation coefficients (the correlations between the neighbouring b.p. steps were omitted from) with and without lag, along with the corresponding lag in ps is presented in the table below.

| **B.p. step shift**<br>System | Maximum correlation without lag | Maximum correlation with lag | Time lag for maximum time-delayed correlation (ps) |
|---|---|---|---|
| WT DNA | -0.2316 | -0.2334 | -1 |
| WT CEBPB | -0.2389 | -0.2029 | -76 |
| WT E2F1-DP1 | -0.2865 | -0.2891 | 648 |
| WT E2F1-DP1-CEBPB | -0.3737 | -0.2687 | -9999 |
| ME DNA | 0.1194 | -0.1143 | 503 |
| ME CEBPB | 0.1745 | 0.1727 | -261 |
| ME E2F1-DP1 | 0.3411 | 0.3452 | 8303 |
| ME E2F1-DP1-CEBPB | 0.3744 | 0.3743 | -55 |

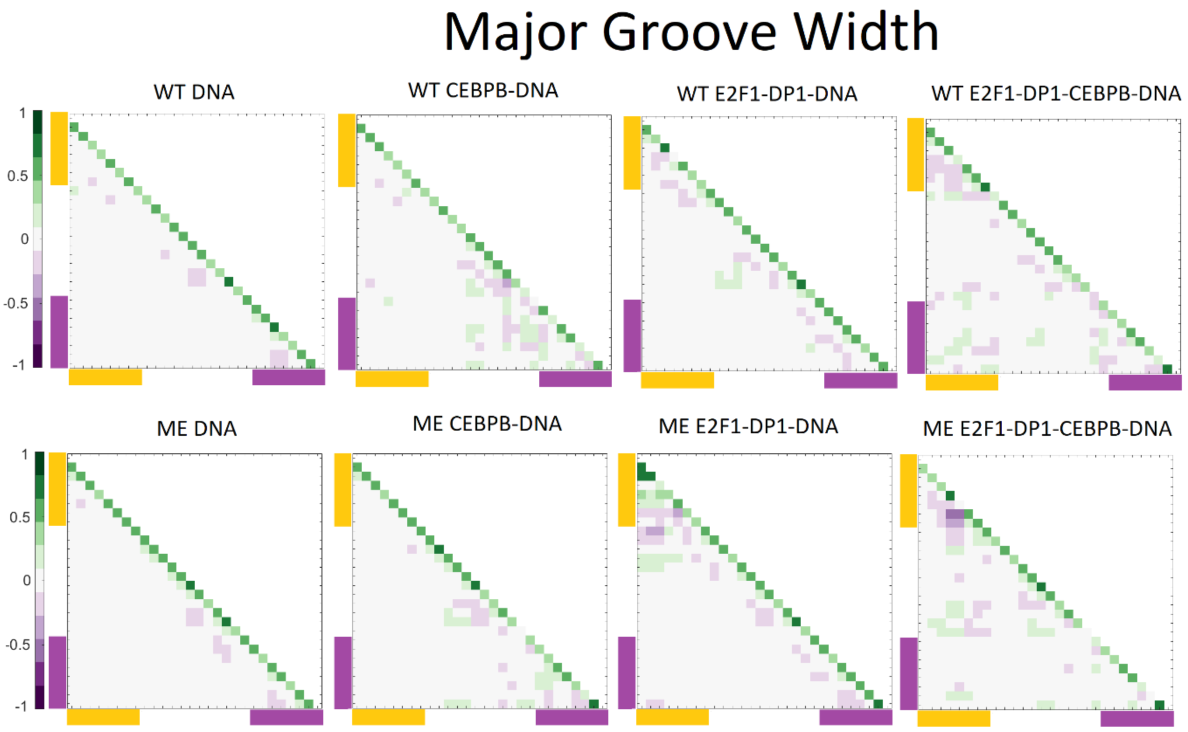


**Figure S27:** Time-delayed correlation coefficient maps for the major groove width, with a maximum allowed time lag of 10 ns along the WT and ME sequences for DNA alone and in complex with either CEBPB/E2F1-DP1 dimers, or with both TFs dimers CEBPB-E2F1-DP1. Correlation coefficients are calculated for the entire (0.1–1.1 μs) trajectories. In all panels, the E2F1-DP1 response element is marked with yellow, the CEBPB response element – with magenta. The maximum correlation coefficients (the correlations between the neighbouring b.p. steps were omitted from) with and without lag, along with the corresponding lag in ps is presented in the table below.

| Major Groove Width System | Maximum correlation without lag | Maximum correlation with lag | Time lag for maximum correlation (ps) |
|---|---|---|---|
| WT DNA | -0.2672 | -0.1737 | -332 |
| WT CEBPB | 0.2583 | -0.2577 | -1 |
| WT E2F1-DP1 | -0.2543 | -0.2369 | 271 |
| WT E2F1-DP1-CEBPB | -0.2792 | 0.267 | 0 |
| ME DNA | -0.2776 | -0.1995 | -511 |
| ME CEBPB | -0.2421 | -0.2095 | -237 |
| ME E2F1-DP1 | -0.3375 | -0.3174 | 89 |
| ME E2F1-DP1-CEBPB | -0.5765 | -0.5687 | -26 |

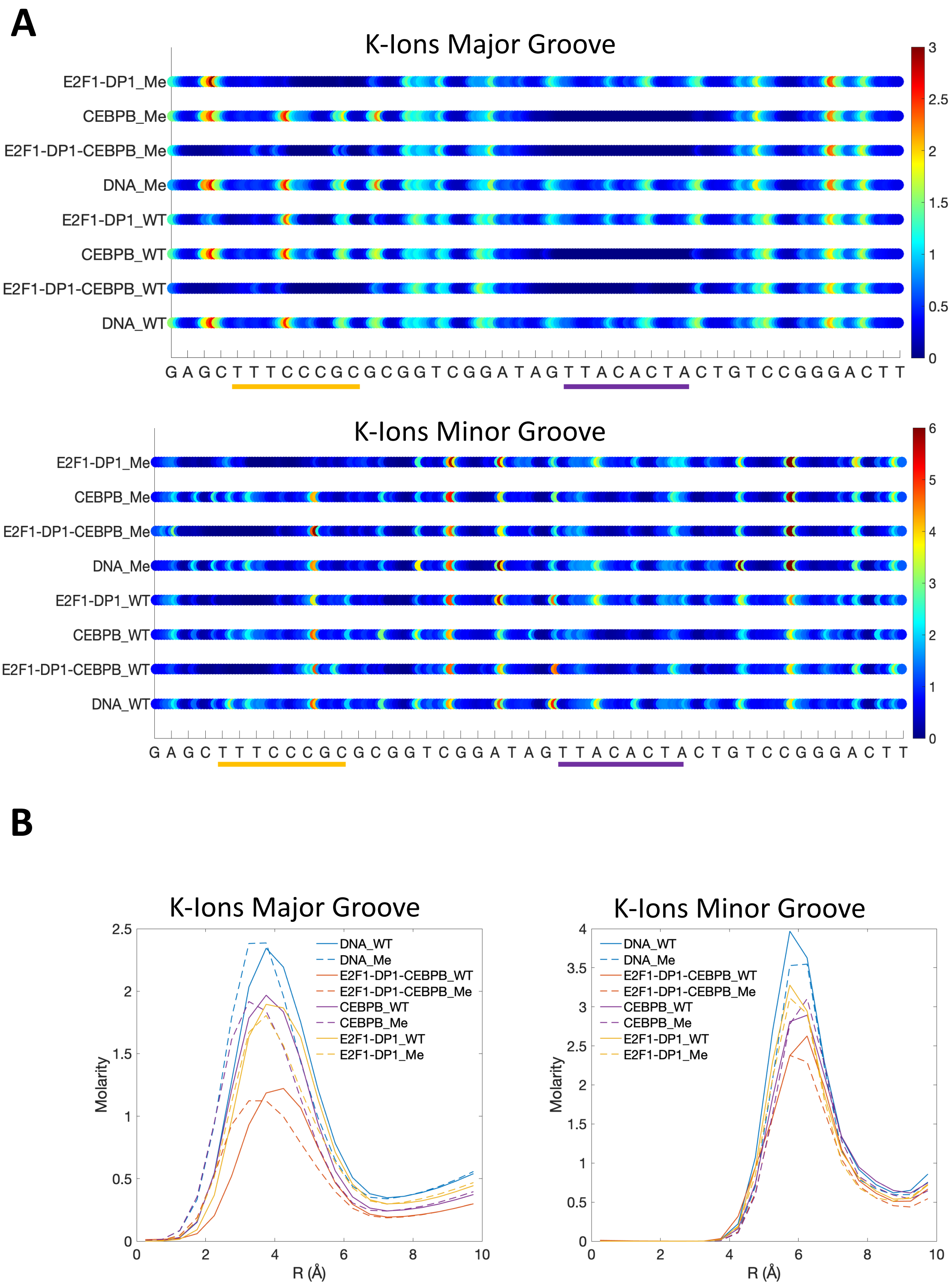


**Figure S28:** K+-molarity in major and minor groove: **A.** along the DNA sequence, **B.** distance from the helical axis.
